# Spectral Matched Filtering in the Butterfly Visuomotor System

**DOI:** 10.1101/2025.06.10.658887

**Authors:** Jack A. Supple, Uroš Cerkvenik, Marko Ilić, Primož Pirih, Aleš Škorjanc, Gregor Belušič, Holger G. Krapp

## Abstract

Color provides an important dimension for object detection and classification. In most animals, color- and motion-vision are largely separated throughout early stages of visual processing. However, accumulating evidence indicates crosstalk between chromatic and achromatic pathways. Here we investigate the spectral sensitivity of the butterfly motion-vision pathway at the level of pre-motor descending neurons (DNs), which connect the brain to thoracic motor centres. Butterflies engage in fast agile flight within often-colorful visual ecologies, which may heighten evolutionary pressure to integrate color- and motion-vision. Indeed, we observed a separation of spectral sensitivities that matches the functional properties of butterfly DNs, such that wide-field optic flow-sensitive DNs involved in stabilisation reflexes have effectively broadband spectral responses, whilst target-selective DNs involved in target-tracking are comparatively narrowband and match conspecific wing coloration. Our findings demonstrate an integration of color- and motion-vision within a pre-motor neuronal bottleneck that controls behavior.

## Introduction

With less than ∼10^6^ neurons^1–3^ and power consumption in the milliwatt range^4, 5^, insects engage with diverse^6^, often agile behaviors^7, 8^, far out-performing modern artificial systems^9^. A foundational principle enabling such high-performance information processing is ‘matched filtering’, whereby sensorimotor systems are specialized to process only task-relevant information^10–13^. Indeed, neurons throughout the insect nervous system are highly task-specialized^11, 13, 14^, with mounting evidence that the entire insect sensorimotor pathway is matched to maximise energy transfer from sensors to motor systems^15–17^. Matched filters are prominent throughout insect visuomotor pathways, ranging from specialisations for visual acuity^11^, spatial gradients of photoreceptor spectral sensitivities^18^, to optic flow-sensitive interneurons that monitor self-motion^12, 16^. However, whilst spectral information provides an important dimension for visual feature extraction^19^, few examples^20^ exist in which color- and motion-vision are integrated into matched-filters spanning both visual dimensions.

Color and motion vision are conspicuously segregated into parallel anatomical pathways in both vertebrates^21^ and invertebrates^22^. In flies, broadband-sensitive photoreceptors R1-6 synapse with lamina monopolar cells (LMCs)^23–25^, which are specialised input channels to downstream motion-detecting neurons^26^. In contrast, photoreceptors R7-8 have differential opsin expression and project long visual fibres bypassing the lamina before terminating in the medulla^25, 27^. Optic lobe output interneurons separate into distinct anatomical tracts^22, 27^, with predominantly achromatic neurons projecting to the lateral protocerebrum and chromatic neurons projecting to the anterior protocerebrum^22, 28^. This separation of parallel chromatic/achromatic-pathways is thought to arise from the trade-off between spectral- and spatiotemporal-acuity, with achromatic vision increasing photon-capture, thereby improving sensitivity and spatiotemporal-resolution^29, 30^. Indeed, color-vision in bees depends on object size, with recognition of small objects depending predominantly on achromatic luminance contrasts, and exclusively chromatic information only used for larger objects^31^. Nonetheless, examples of crosstalk between chromatic/achromatic pathways are emerging^20, 32–35^. For example, asymmetric spectral-sensitivities of ON/OFF motion-sensitive T4/T5 cells (the first stage of motion detection^36^) in *Drosophila* enhances the detection of approaching colored objects, demonstrating potential advantages of integrated chromatic motion-vision^20^.

Descending neurons (DNs) occupy a focal point in the insect nervous system by connecting the brain to thoracic motor centres^37, 38^. As an information bottleneck, DNs integrate sensory modalities to improve the speed and precision of sensorimotor control^39, 40^, and subpopulations of specialised DNs are matched to control specific aspects of behavior^38^. Notably, stabilisation reflexes and goal-directed behaviors such as target-tracking are largely designated to separate DN populations^41^, enabling specialisation for the different dynamics of each task^42^. DNs sensitive to wide-field optic flow (WFDNs) have been described in a variety of insects^43–55^, and respond selectively to a preferred pattern of wide-field motion across a large visual area^43^. WFDNs receive input from optic flow-sensitive output neurons from the optic lobe, project from the lateral protocerebrum to the thoracic ganglia, and are thought to coordinate optomotor stabilization reflexes^43, 44, 54, 56, 57^. In contrast, target-selective DNs (TSDNs) are sensitive to small objects moving in a preferred direction^41, 47, 50^, featuring localized receptive fields aligned with other neuroanatomical specialisations for target-tracking such as increased sensitivity and spatiotemporal acuity^11, 58, 59^. For example, TSDNs in predatory Odonates and Dipterans align with visual acute zones upon which prey is fixated during interception^40, 58–61^, whilst TSDNs in non-predatory Dipterans sample from the ‘love-spot’^62^ acute zone to track conspecifics^47, 63, 64^. TSDNs are thought to receive input from small-target movement-detector interneurons projecting from the lobula complex^65, 66^ and/or medulla^63, 67, 68^, and descend to innervate thoracic motor centers^47, 59, 63^, often directly synapsing with motor neurons^47, 63^.

As a sensorimotor interface, DNs present an interesting stage at which to investigate the integration of color- and motion-vision. Both WFDNs and TSDNs respond to visual inputs, even in immobilized animals^47, 50, 51^, suggesting that additional dendritic gating mechanisms and/or parallel pathways of DNs are integrated to coordinate behavior. It is presently unknown whether motion-sensitive DNs are achromatic and gated by chromatic information depending on behavioral context, or whether motion-sensitive DNs are inherently spectrally selective. In dragonflies, TSDNs are UV/blue-sensitive^69^, matching the spectral sensitivity of the dorsal retina^70, 71^. However, dragonflies contrast prey silhouettes against the sky by hunting from below^72–74^, so this short-wavelength specialisation increases the signal-to-noise ratio of extremely small (<1°) targets^72, 75, 76^. Similarly, WFDNs^53, 69, 77^ and their inputs from the lobula complex^78–81^ are driven primarily by green-sensitive photoreceptors in most tested insects, matching the spectral-sensitivity of optomotor behaviors^80–93^. Optomotor green-sensitivity is presumed to improve photon-catch by matching the predominantly green diurnal terrestrial background^90, 93^, with optomotor responses of crepuscular species correspondingly blue-shifted^93^. Additional photoreceptor inputs nonetheless improve wide-field motion detection in flies^32^ and bees^77^ by increasing photon-catch across the visible spectrum^32^.

We investigated the integration of color and motion-vision at the level of DNs in butterflies, a diverse clade^94^ remarkable for both their highly manoeuvrable flight^95^, and their often-colorful visual ecologies^96^. Butterflies frequently engage in fast aerial pursuits during both courtship and territorial conflicts^97–100^. Males competing for territory execute fast spiralling flights, with both males sequentially exchanging status as chaser and evader until one concedes^97, 100–103^. Similarly, courtship in many species is initiated by males pursuing flying conspecific females^98, 103–105^. Both sexes broadly assess mate quality from multiple cues including color^106–109^, size^107, 109^, and odor^106–108^, with females suggested to use aerial pursuit to identify fitter mates^109, 110^. The flight performance demanded by sexual behaviors necessitates a combination of robust stabilisation reflexes functioning in parallel with a specialised target tracking system^14, 42, 111–113^. The ubiquitous use of color throughout butterfly behavioral ecology may impose selective evolutionary pressures to integrate color- and motion-vision. Indeed, butterfly wide-field motion-vision is sensitive to chromatic-only contrasts^35^, unlike in flies^92^ and bees^85^.

Butterfly color vision is supported by nine photoreceptors R1-9 per ommatidium, forming a fused and tiered rhabdom ^114^, with diverse patterns of opsin and pigment expression across the retina depending on species^115^. R1&2 (and in some species R9) project long visual fibres to the medulla, whilst R3-8 are short visual fibres, terminating in the lamina^116, 117^. Inter-photoreceptor synapses produce additional combinations of spectral opponency^116, 118^, and an array of spectrally-narrowband interneurons span from 300-650nm in the mushroom bodies^119–121^. Whilst most LMCs are spectrally-broadband, receiving multiple photoreceptor inputs^93, 116, 122^, some heterogeneity between LMC types suggests the potential for spectrally-selective motion-vision^122^. Indeed, one early study reported species-dependent spectral-selectivity in butterfly WFDNs^53^, and motion-sensitive feedback neurons projecting from the protocerebrum to the medulla are sensitive to chromatic-only contrasts^34^. In this study we report the functional properties of a subset of butterfly motion-sensitive DNs, finding WFDNs and TSDNs akin to those reported in other species. We then compared DN spectral sensitivities, finding a functional separation whereby WFDNs have effectively broadband spectral responses matching optomotor reflexes, whilst TSDNs are comparatively narrowband and match conspecific wing coloration.

## Results

### Diversity of Butterfly Wing Coloration and Retinal Composition

We compare two species of New World Nymphalid butterflies: the Monarch butterfly *Danaus plexippus*, and the Blue-wave butterfly *Myscelia cyaniris* (referred to hereon by their genus). With a last common ancestor ∼78.85 MYA^94^, *Danaus* and *Myscelia* have notable divergence in wing coloration (Fig. 1A). *Danaus* are aposematic, signalling their toxicity to predators via a distinctively bright-orange wing coloration^124, 125^ (Fig. 1Ai). Reflectance increases between 500-700nm, inflecting at ∼540nm (Fig. 1Bi). Whilst coloration is slightly red-shifted in males and migration-stage butterflies^126^, sexual dimorphism is minimal, limited to enlarged black vein widths in females, and two black androchonial pheromone glands on the male hindwings (Fig. 1Ai). *Myscelia* sexual dimorphism is more pronounced (Fig. 1Aii). Both sexes are striated with light bands across both wings (Fig. 1Aii). However, males possess a vibrant blue iridescence, with a reflectance maximum at ∼370nm and half-max at 460nm (Fig. 1Bii). This coloration is largely absent in females, although the light stripes possess a subtle shade of violet (Fig. 1Aii).

**Figure 1:**
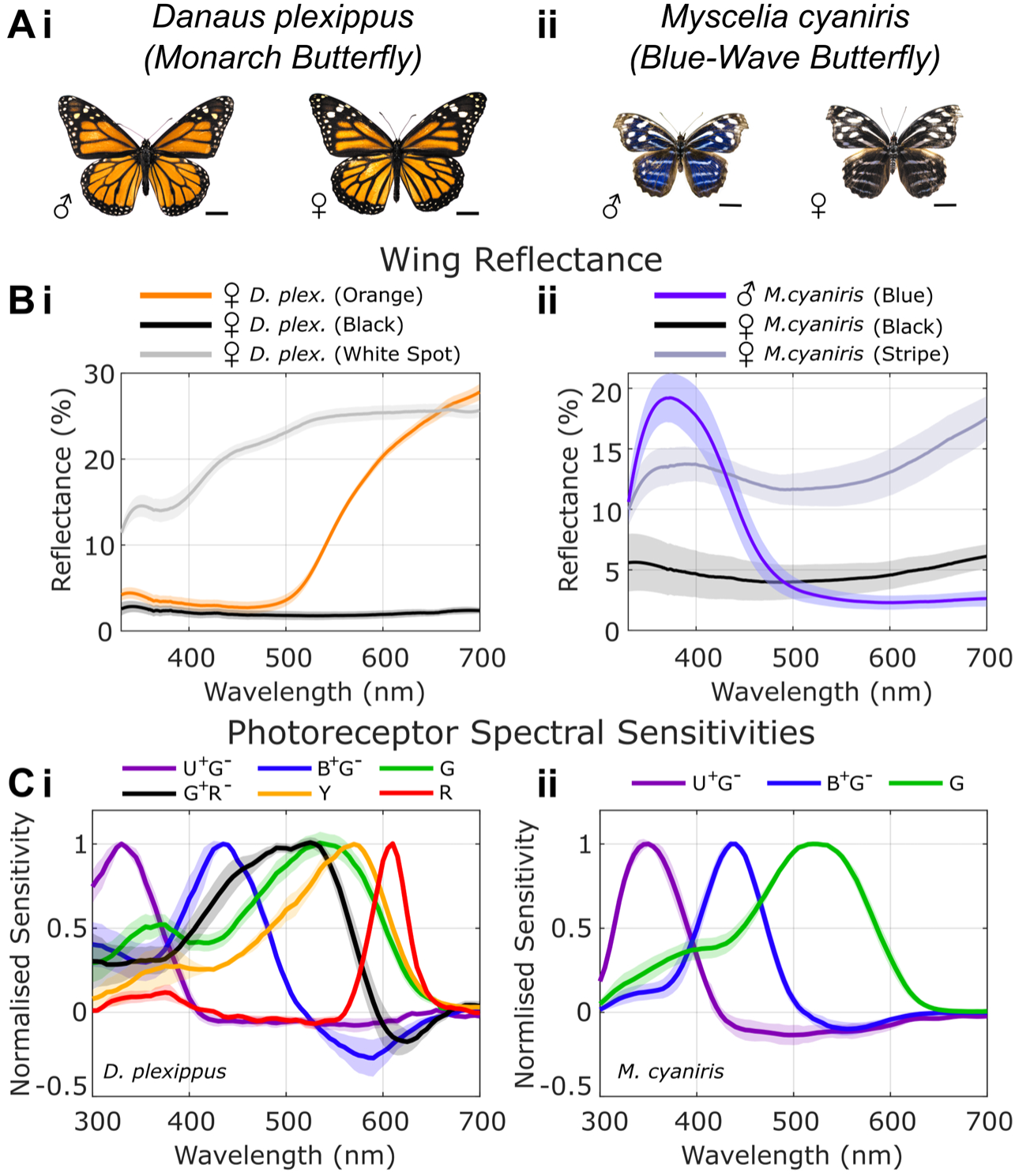
Photoreceptor spectral sensitivities and wing coloration in *D. plexippus* and *M. cyaniris*. (A) Male and female specimens of (i) *D. plexippus* and (ii) *M. cyaniris*. Scale bars 1 cm. All specimens displayed at the same scale for comparison. (B) Wing reflectance measurements. (i) *D. plexippus* forewing reflectance from orange, and black perimeter regions, and white spots. N=2 animals. (ii) *M. cyaniris* reflectance from male blue forewing (N=3 animals), female black regions (combined fore- and hindwings, N=2 animals), and female white/violet stripes (combined fore- and hindwings, N=2 animals). (C) Photoreceptor spectral sensitivities from (i) *D. plexippus* (N = 4/5/8/3/6/2, U^+^G^-^/B^+^G^-^ /G/G^+^R^-^/Y/R cells from 4 animals), and (ii) *M. cyaniris* (N=4/4/7, U^+^G^-^/B^+^G^-^/G cells from 2 animals).

*Danaus* and *Myscelia* are also divergent in their retinal composition (Fig. 1C). Previous work demonstrated that Nymphalid photoreceptor rhodopsin expression patterns produce either simple or complex retina types depending on sex and species^118, 127^. The basic (simple) composition of Nymphalid retinas includes R3-8 expressing long-wavelength, green- or yellow-peaking rhodopsins, whilst R1-2 express either UV or blue-peaking rhodopsins^118, 127^. *Myscelia* retinas correspond to this simple design, with three photoreceptor types peaking in UV, blue, and green (Fig. Cii), with inhibitory green-opponency in UV and blue photoreceptors (U^+^G^-^ and B^+^G^-^, respectively). Complex retinas contain all ommatidial types found in simple retinas; however, an additional ommatidial type expresses a long-wavelength (green) rhodopsin in R1-2^127–129^. These ommatidia also contain a red screening pigment that filters light reaching the photoreceptors at the base of the retina: the proximal R5-8 are therefore yellow-shifted (termed ‘Y’ receptors), while the basal R9 is red-sensitive^127, 129^. Inhibitory synapses between red-sensitive R9 and green R1-2 produces green-red opponent (G^+^R^-^) cells^129^. The retina of *Danaus* corresponds to this complex form, with red-sensitive R9, yellow-sensitive R5-8, and G^+^R^-^ extending spectral-sensitivity to 650nm (Fig. 1Ci).

### Parallel pathways of Motion-Sensitive Descending Neurons in Butterflies

To investigate the spectral sensitivity of butterfly DNs, we presented moving patterns in the frontal visual field using a customised display system comprised of a multi-LED galvo scanner and an RGB DLP projector (Fig. S1). Using the DLP, we first classified DNs into two functional categories based on their sensitivity to different moving patterns (Fig. 2). Wide-field optic flow-sensitive descending neurons (WFDNs) were classified as neurons with (i) directionally selective responses to wide-field moving gratings with approximately cosine tuning curves (Fig. 2C-D, row i); and (ii) monotonically increasing responses to increasing target sizes (Fig. 2C-D, row ii). Target-selective descending neurons (TSDNs) were defined as neurons that (i) responded weakly (<20 spikes/s) to wide-field moving gratings; (ii) responded to targets with a distinct preferred direction of motion; and (iii) displayed non-linear target size tuning curves with a single distinct peak^50^.

**Figure 2:**
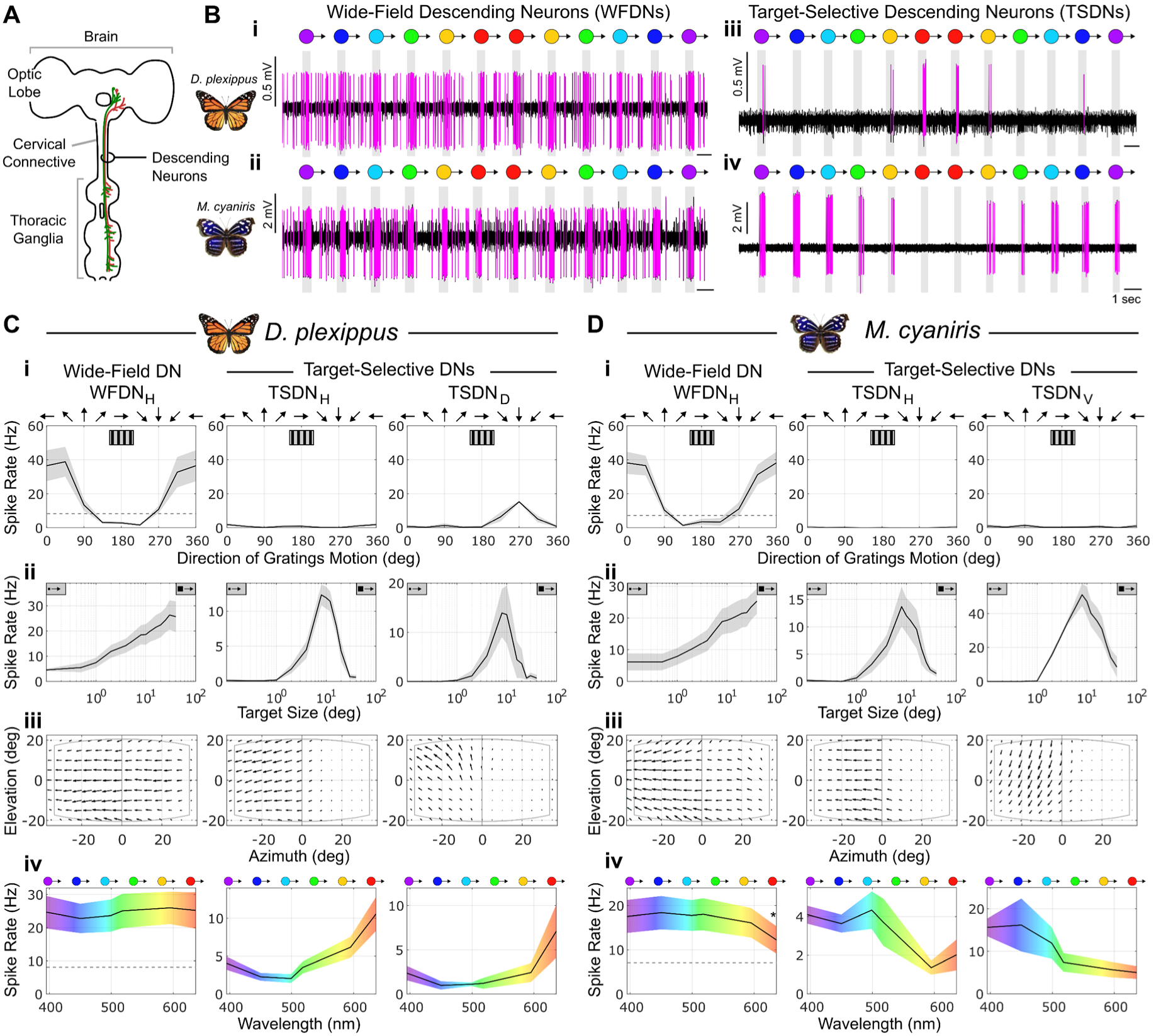
Spectral sensitivities of Wide-Field and Target-Selective Descending Neurons in *D. plexippus* and *M. cyaniris*. (A) Schematic of butterfly nervous system and descending neurons (DNs). DNs were recorded from the cervical connective. (B) Example raw extracellular voltage traces of (i-ii) WFDNs and (iii-iv) TSDNs in *D. plexippus* (top row) and *M. cyaniris* (bottom row) responding to 8°-diameter circular bright objects of different color moving in the preferred direction. (C-D) Most-frequently encountered WFDNs and TSDNs in (C) *D. plexippus* (WFDN_H_, N=7 cells from 6 animals; TSDN_H_, N=11 cells from 9 animals; TSDN_D_, N=3 cells from 2 animals), and (D) *M. cyaniris* (WFDN_H_, N=8 cells from 5 animals; TSDN_H_, N=5 cells from 4 animals; TSDN_V_, N=6 cells from 5 animals). (i) Responses to gratings moving in different directions in the frontal visual field. Dashed-grey line shows average spontaneous firing rate. (ii) Responses to dark objects of different side-length subtended angles (degrees) moving in the preferred direction through the receptive field. (iii) DN receptive fields. Vector directions and magnitudes represent the local preferred direction and normalised motion sensitivity of the neuron across the visual field. (iv) Spectral tuning curves showing DN responses to a bright 8°-diameter circular object moving at a constant velocity of 115 °/sec. Dashed-grey line shows WFDN spontaneous firings rates. WFDN responses statistically significantly different as determined from ANOVA (p<0.01) followed by post-hoc t-test comparisons indicated (*, p<0.05).

In both species, the most frequently encountered WFDN was sensitive to horizontal motion (WFDN_H_; Fig. 2C-D, left), with a baseline spontaneous firing rate of 8.1±6.1 and 7.1±6.0 spikes/sec (mean±SEM; *Danaus* and *Myscelia*, respectively), increasing to a maximum of 38.8±8.7 and 38.1±6.5 spikes/sec in the preferred direction (towards the ipsilateral side), and decreasing to 1.6±0.47 and 1.4±0.6 spikes/sec in the anti-preferred direction (*Danaus:* N=7 cells, 6 animals*; Myscelia:* N=8 cells, 5 animals; Fig. Ci-Di). WFDN_H_ receptive fields were binocular in both species (Fig. Ciii-Diii), although *Myscelia* WFDN_H_ was more lateralised to the ipsilateral visual field (Fig. Diii).

The most frequently encountered TSDNs in *Danaus* were sensitive to either horizontal or ventral-dorsal target motion (TSDN_H_ and TSDN_D_; Fig. C middle/right columns, respectively; TSDN_H_: N=11 cells, 9 animals, and TSDN_D_: N=3 cells, 2 animals). The most frequently encountered TSDNs in *Myscelia* were sensitive to either horizontal or dorsal-ventral target motion (TSDN_H_ and TSDN_V_, Fig. D middle/right columns, respectively; TSDN_H_: N=5 cells, 4 animals; TSDN_V_, N=6 cells, 5 animals). All measured TSDNs in both species had negligible spontaneous firing rates (<1 spikes/sec), were maximally sensitive to a target size of 8° (dark contrast, Fig. 2C-Dii), and had ipsilateral receptive fields (Fig. 2C-Diii). TSDN responses to moving gratings were absent, except for *Danaus* TSDN_D_, which responded to gratings moving in the same preferred direction as for target responses (Fig. 2Ci, right column), albeit with a lower amplitude than WFDN_H_.

### Functional Separation of Butterfly Descending Neuron Spectral Sensitivities

WFDN and TSDN spectral tuning curves were measured using the multi-LED galvo scanner, which presented moving bright objects subtending 8° diameter. WFDN_H_ spectral sensitivities were broadband in both species (Fig. 2C-Div). However, *Danaus* WFDN_H_ displayed a subtle reduction in sensitivity for blue objects (reduction of normalised mean spike rate by -7%; ANOVA p=0.02; t-test comparison to average, p=0.04 without adjusting for multiple comparisons; Fig. 2Civ), whilst *Myscelia* WFDN_H_ had a more pronounced reduction in sensitivity for red objects (reduction of normalised mean spike rate by -38%; ANOVA p<10^-7^; t-test comparison to average, p<10^-3^ with Bonferroni correction; Fig. 2Div). *Myscelia* WFDN_H_ responses were well-explained by green photoreceptor input (R^2^=0.70), with additional B^+^G^-^-receptor input improving the modelled response (R^2^=0.85; green:blue=1:0.37; Fig. S3). *Danaus* WFDN_H_ responses were poorly modelled by green-only photoreceptor input (R^2^<0: i.e. best-fit responses worse than a straight line; Fig. S3C) but were compatible with a range of multi-photoreceptor input combinations (R^2^>0.9; Fig. S3D).

In contrast, TSDNs in both species had pronounced narrowband spectral-sensitivity (Fig. C-Div). In *Danaus*, TSDN_H_ and TSDN_D_ were maximally sensitive to red (636nm), and minimal for green (499nm), with a smaller secondary maximum for UV (395nm). In *Myscelia*, TSDN_H_ and TSDN_V_ were maximally sensitive to short wavelengths (TSDN_H_ maximum=470nm; TSDN_V_ maximum=420nm; based on smoothened estimates, Fig. S9), and were minimally sensitive to orange/red (Fig. 2Div). Best-fit linear combinations of photoreceptor inputs to *Danaus* TSDNs both had Y-receptor as a primary excitatory input (Fig. S3E-F), but were also compatible with combinations of inhibitory G^+^R^-^/Y/R-receptor inputs onto green-receptor excitation (R2>0.9; Fig. S3E-F). While *Myscelia* TSDN_H_ responses were poorly modelled by linear combinations of photoreceptor inputs (R^2^=0.5, Fig. S3K), *Myscelia* TSDN_V_ was compatible with predominantly B^+^G^-^-photoreceptor input (R^2^=0.86; Fig. S3L).

Whilst these TSDNs were the most frequently encountered during experiments, other TSDNs were also found less regularly, including TSDN_V_ in *Danaus*, which displayed similar functional properties to *Myscelia* TSDN_V_, but shared the UV/red spectral-sensitivity of *Danaus* TSDN_H_ and TSDN_D_ (Fig. S4A). Although we focussed on measuring male *Myscelia* TSDNs, we found no difference in TSDN spectral sensitivities between male and female *Danaus* TSDN_H_ (Fig. S4B).

### Descending Neuron Spectral Sensitivities are Contrast Dependent

To further investigate DN spectral sensitivities, we tested spectral responses to an extended sequence of moving patterns using the RGB DLP (Fig. 3&4). In both species, we found no statistically significant differences in WFDN_H_ response gain to moving gratings with different colors (Fig. 3A; non-parametric permutation tests, p>0.05 after multiple test corrections; see Fig. S5A). Similarly, there were no statistically significant differences in WFDN_H_ responses to dark objects of different sizes (Fig. 3Bi-ii) or different velocities (Fig. 3Ci-ii) when moving on a bright background (non-parametric permutation tests, p>0.05 after multiple test corrections, Fig. S5B,D). However, for bright objects moving on a dark background, there was a subtle reduction in *Danaus* WFDN_H_ responses for blue targets, and a reduction in *Myscelia* WFDN_H_ responses for red targets, for variations in both target size (Fig. 3Biii-iv) and velocity (Fig. 3Ciii-iv). These response reductions matched those observed for the multi-LED galvo stimuli (Fig. 2C-Div) but were only found to be statistically significant for *Danaus* WFDN_H_ (Fig. 3C-Diii; non-parametric permutation tests, p≤0.05 after multiple test corrections, N=7 cells, see Fig. S5C,E for statistical analyses).

**Figure 3:**
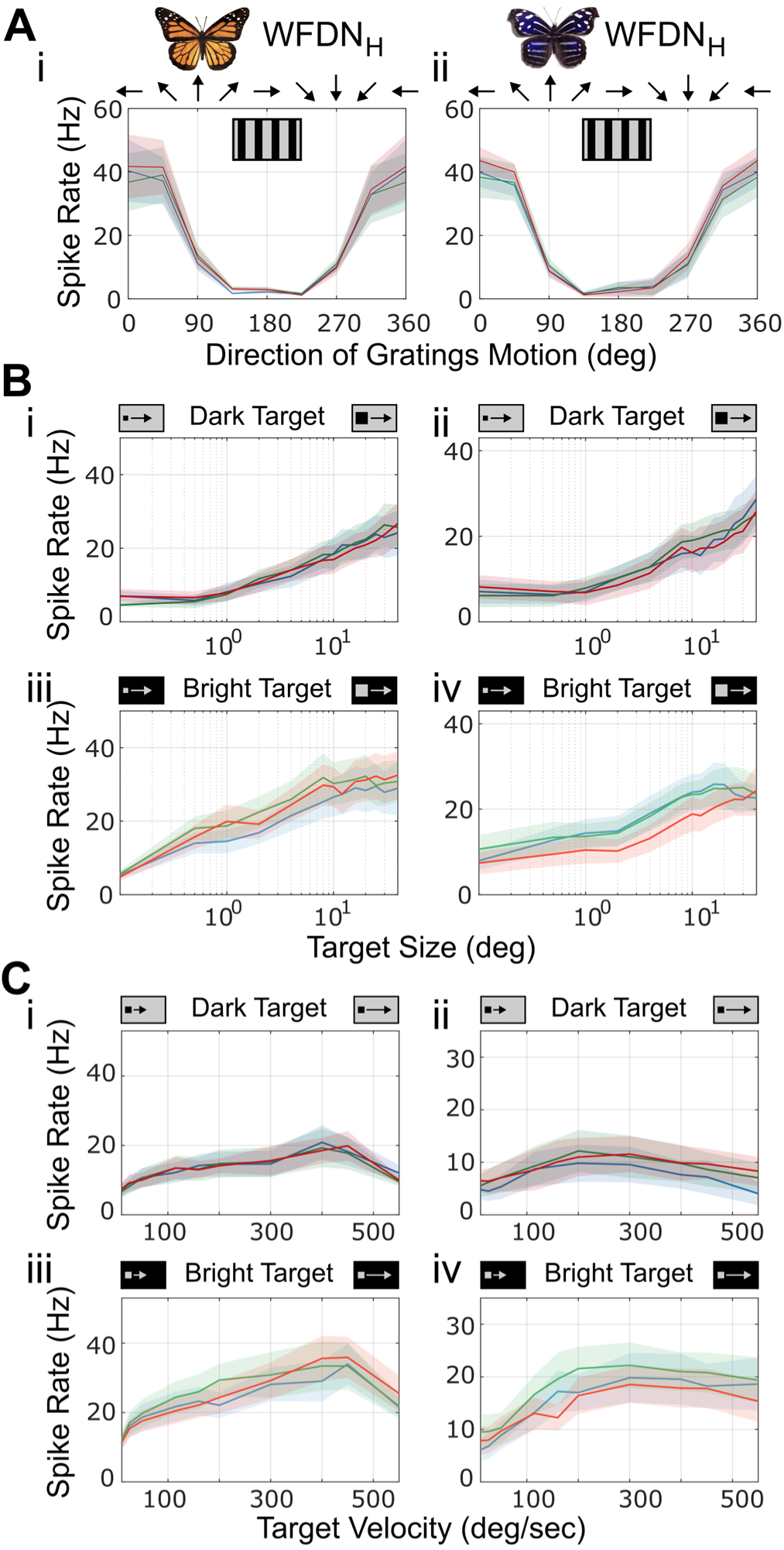
Wide-Field Descending Neurons (WFDNs) display subtle contrast-dependent spectral selectivity. **(A)** WFDN_H_ responses to gratings moving along different directions in the frontal visual field (10° wavelength, 5 Hz temporal frequency) for (i) *D. plexippus,* and (ii) *M. cyaniris*. **(B)** Size tuning curves for (i-ii) dark and (iii-iv) bright square objects moving in the preferred direction at constant 115 °/sec velocity. **(C)** Velocity tuning curves for (i-ii) dark and (iii-iv) bright square objects with constant side lengths subtending 4° and moving in the preferred direction. *D. plexippus* WFDN_H_ (left column, N=7 cells, 6 animals). *M. cyaniris* WFDN_H_ (right column, N=8 cells from 5 animals). See Figure S4 for statistical analysis. Note B-C share y-axis scale for comparison within species.

Unlike WFDN_H_, TSDNs in both species displayed distinct spectral selectivity across a range of target sizes and velocities (Fig. 4, Fig. S6). Responses to dark targets moving on a bright background were similar in magnitude for red, green, and blue light, although both species had a slight reduction in responses to blue targets on dark backgrounds for variations in both target size (Fig. 4Ai-ii) and velocity (Fig. 4Bi-ii). Spectral selectivity was more prominent for bright targets moving on a dark background, consistent with measurements using the multi-LED galvo system (Fig. 2C-Div). In *Danaus*, TSDN_H_ displayed statistically significantly larger peak firing rates for bright red targets (Fig. 4Aiii; non-parametric permutation tests p<0.05 after multiple test corrections, N=10 cells), whilst *Myscelia* TSDN_V_ responded more to bright blue targets (Fig. 4Aiv; non-parametric permutation tests p<0.05 after multiple test corrections, N=6 cells). These trends were consistent for variations in both target size (Fig.4Aiii-iv) and velocity (Fig.4Biii-iv; see Fig. S6 for statistical analyses).

**Figure 4:**
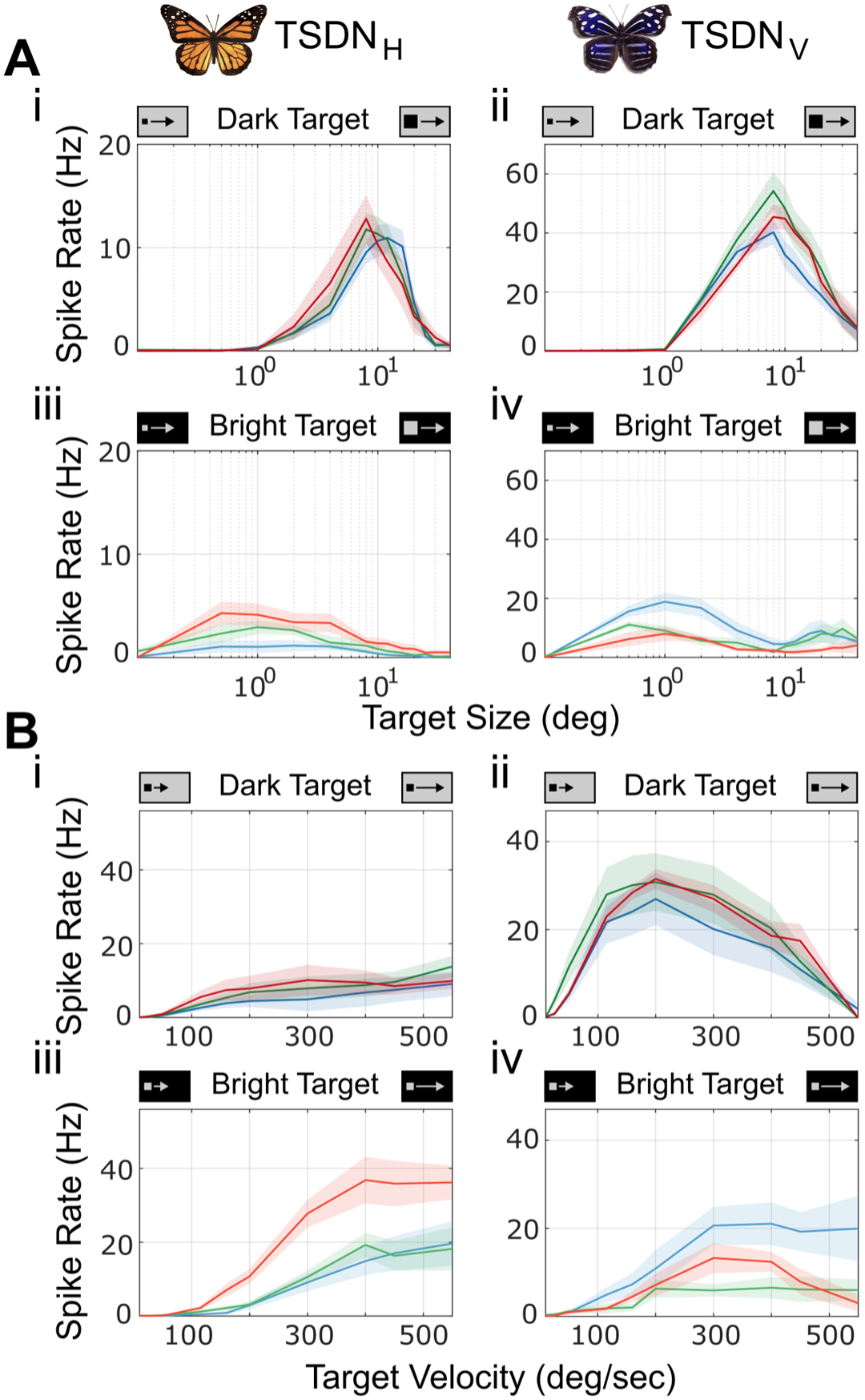
Butterfly TSDN spectral sensitivity depends on target size, velocity, and contrast. **(A)** TSDN size tuning curves for (i-ii) dark and (iii-iv) bright square objects moving in the preferred direction at constant 115 °/sec velocity. **(B)** TSDN velocity tuning curves for (i-ii) dark and (iii-iv) bright square objects with constant side lengths subtending 4° and moving in the preferred direction. *D. plexippus* TSDN_H_ (left column, N=10 cells from 8 animals). *M. cyaniris* TSDN_V_ (right column, N=6 cells from 6 animals). See Figure S5 for statistical analysis. Note A-B share y-axis scale for comparison within species.

Notably, we found that *Danaus* TSDNs responded with highest peak firing rates to bright red targets moving at velocities >400°/sec (Fig. 4Biii). These fast moving bright red targets acted as a super-stimulus, eliciting ∼1.9x the response (maximum of 36.8±18.9 spikes/sec at 400°/sec) compared to any other tested stimulus parameter (19.7±18.2 spikes/sec for bright blue targets at 550 °/sec). We confirmed that this high-velocity sensitivity was directionally selective (Fig. S4E). We also observed substantial differences in TSDN size preference for dark vs bright moving targets (Fig. 4A). In both species, dark objects produced peak responses for ∼8° target diameters (Fig. 4Ai-ii), whilst bright targets produced peak responses at diameters of ∼1° (Fig. 4Aiii-iv).

### Wide-Field Optic Flow-Sensitive Descending Neuron (WFDN) Spectral Sensitivities Match Optomotor Behavior

To test whether DN spectral sensitivities match behavior, we first measured the spectral sensitivity of stabilising optomotor head movements in both species (Fig. 5, Fig. S7). Perched butterflies rotated their head yaw orientation in response to gratings moving horizontally on a spectrally-calibrated computer monitor (Movies S1-S2). The amplitude of head yaw rotation was maximal at a grating temporal frequency of ∼20 cycles/sec in both species (Fig. 5B-D). When tested with differently colored gratings, *Danaus* exhibited a reduction in maximal yaw amplitude for blue gratings (Fig. 5Di-E; 7.3±1.6, 8.8±1.2, and 8.4±1.1 degrees, blue/green/red respectively. N=17 animals. Repeated-measures ANOVA, p<0.01, and post-hoc pairwise t-tests with Bonferroni correction, blue-green: p<0.01; blue-red: p=0.07; green-red: p>0.5). In contrast, *Myscelia* head yaw rotation amplitudes were maximal for green, and minimal for red gratings (Fig. 5Dii,E; 6.6±1.3, 7.1±1.2, and 5.1±1.4 degrees, blue/green/red, respectively. N=8 animals. Repeated-measures ANOVA, p=0.01, and post-hoc pairwise t-tests with Bonferroni correction, blue-green: p>0.5; blue-red: p=0.08; green-red: p=0.01. See Fig. S7 for raw data). This reduction in optomotor yaw amplitude for blue gratings in *Danaus*, and red gratings in *Myscelia* matches the spectral sensitivity of WFDNs (Fig. 2-4).

**Figure 5:**
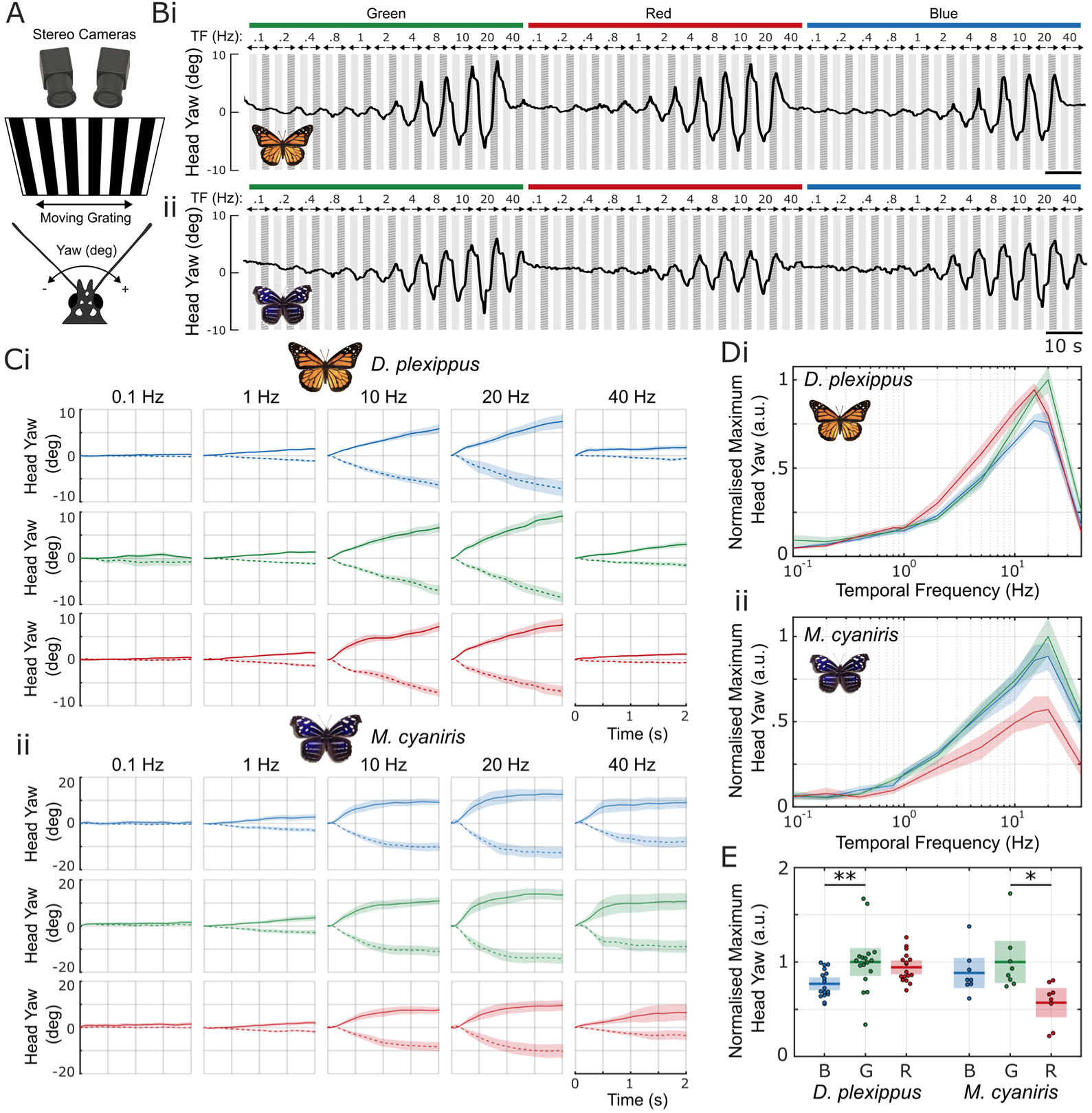
Spectral Sensitivity of Butterfly Optomotor Head Movements. **(A)** Perched butterflies rotated their head in response to gratings (20° spatial wavelength) moving horizontally on a spectrally calibrated computer monitor. 3D head rotations were reconstructed from stereo-videography at 250 fps. **(B)** Example head yaw response from (i) *D. plexippus* and (ii) *M. cyaniris* to red, green, or blue gratings moving at temporal frequencies (TFs) ranging from 0.1 to 40 Hz. Gratings moving left and right indicated by light and dark grey periods, respectively. **(C)** Time-course of head yaw rotations (mean ± SEM) for (i) *D. plexippus* (N = 17 animals) and (ii) *M. cyaniris* (N = 8 animals). Dashed and solid lines correspond to head responses to gratings moving left (i.e. in the animal’s negative yaw direction) and right, respectively. Data displayed relative to head yaw position at grating movement onset. Only 5 of the 11 tested temporal frequencies are shown. **(D)** Normalised maximum of averaged left and right head yaw amplitudes for each tested temporal frequency for (i) *D. plexippus* (N = 17 animals) and (ii) *M. cyaniris* (N = 8 animals). Temporal frequency tuning curves are first normalised to the overall mean for each animal, then to the maximum of means across each color condition (i.e., green). Data show mean ± SEM. **(E)** Comparison of normalised maximum head yaw amplitudes for each grating color. Peak head yaw responses for different grating colors were statistically significantly different for both *D. plexippus* and *M. cyaniris* (repeated-measures ANOVA, p≤0.01). Post-hoc Bonferroni-corrected pairwise comparisons indicated as * for p≤0.05, and ** for p≤0.01. *D. plexippus*: blue-green: p<0.01; blue-red: p=0.07; green-red: p>0.5. *M. cyaniris*: blue-green: p>0.5; blue-red: p=0.08; green-red: p=0.01. See Figure S6 for non-normalised raw data comparisons.

To test whether TSDN spectral sensitivities match target-tracking behaviors, we extended our optomotor head rotation protocol to include bright targets of varying size moving horizontally across the screen (Fig. S8). We did not observe any head rotations to moving targets in *Myscelia*; however, *Danaus* occasionally rotated their head to follow target movement (Fig. S8A). Whilst there was a subtle increase in head yaw response amplitudes for red targets and a reduction for small blue targets, this trend was statistically insignificant after accounting for multiple testing (p>0.05, N=20 animals. Fig. S8C-H). We therefore proceeded to investigate putative functional relationships between TSDN spectral sensitivity and goal-directed behavior by comparing TSDN spectral alignment with conspecific wing reflectance.

### Butterfly TSDN Spectral Tuning Matches Conspecific Wing Reflections under Natural Skylight

To test whether butterfly TSDN spectral sensitivities (Fig. 2C-D) match conspecific wing reflectance (Fig. 1B), we calculated spectral alignment as the cosine similarity between each spectral function (equivalent to the Pearson correlation coefficient for two vectors; Fig. 6). In *Danaus*, TSDN spectral tuning was closely aligned to wing reflectance (0.95 for TSDN_H_ and 0.82 for TSDN_D_; Fig. 6C). The near-perfect matching between *Danaus* TSDN_H_ and wing reflectance was impeded by the increase in UV-blue sensitivity between 395-450nm; indeed, the alignment between TSDN_H_ and wing reflectance increased to 0.97 in the 450-650nm band. TSDN_D_ alignment in the 450-650nm band increased to 0.86 but remained less than that of TSDN_H_ due to the slightly red-shifted TSDN_D_ sensitivity (TSDN_H_ half-max = 586nm; TSDN_D_ half-max = 608nm; orange wing reflectance half-max = 566). In *Myscelia*, wing reflectance spectra were less closely aligned to TSDN spectral sensitivities (0.60 for TSDN_H_ and 0.85 for TSDN_V_; Fig. 6C), attributable to the rightward shift in TSDN spectral maxima (TSDN_H_ maximum: 470nm; TSDN_V_ maximum: 420nm) from the maximum wing reflectance (maximum: 370nm).

**Figure 6:**
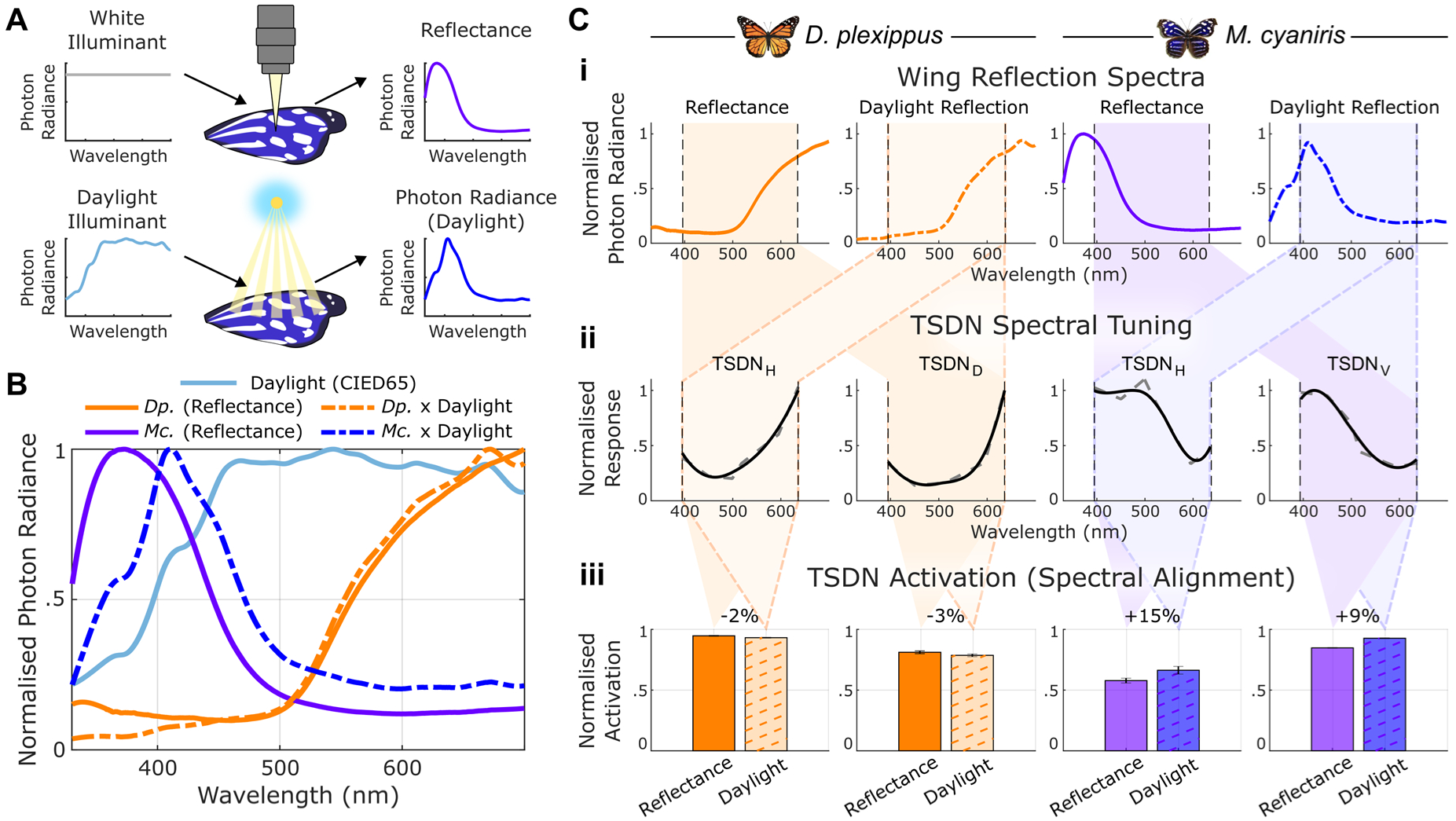
Butterfly TSDN Spectral Tuning Matches Conspecific Wing Reflections under Daylight Illuminant. **(A)** Wing reflectance is a standardised metric referring to the fraction of illuminant photon radiance reflected as a function of wavelength. The normalised (i.e., unitless) wing reflectance spectrum therefore equals the normalised reflected photon radiance under a calibrated white illuminant. Under natural conditions, however, the spectral content of reflected photon radiance is a combination of both the wing reflectance and the illuminant (e.g. daylight). **(B)** Comparison of normalised wing reflectance and estimates of natural wing reflection under daylight, calculated by multiplying wing reflectance with the standard average daylight illuminant, CIED65. *Dp.*: *D. plexippus*; *Mc.*: *M. cyaniris*. **(C)** Calculation of spectral alignment between wing reflectance and reflected daylight photon radiance (top row) and TSDN spectral tuning (middle row). Raw average TSDN tuning curve (dashed grey) shown with smoothed TSDN tuning curve (black). Spectral alignment is calculated as the cosine similarity across the 395-636 nm range (i.e. the range used for TSDN measurements; bottom row). Percentage changes in spectral alignment between standard wing reflectance (solid color) and daylight wing reflection (dashed) are displayed. Error bars represent standard deviation of alignment estimates for different levels of TSDN tuning curve smoothing (see Figure S9).

However, our measurements of wing reflectance spectra were acquired using calibrated white light, producing the standardised reflectance (Fig. 6A). In nature, the reflected photon radiance depends on both the wing reflectance and the spectral content of the natural illuminant (usually skylight; Fig. 6A)^130^, which is attenuated below 450nm (Fig. 6B)^131, 132^. To simulate reflected photon radiance under natural conditions, we multiplied the wing reflectance with the standard daylight illuminant, CIED65 (Fig. 6B). The reflected photon radiance of *Danaus* wings closely resembles reflectance due to the relatively flat skylight spectrum above 450nm (Fig. 6B), with a slight reduction in spectral alignment of ∼2% to 0.94 in TSDN_H_, and ∼3% to 0.81 in TSDN_D_ (Fig. 6C, S9). However, due to skylight short-wavelength attenuation, the natural reflected photon radiance maximum of male *Myscelia* wings shifted from 370nm to 410nm (Fig. 6B). This rightward shift increases alignment with *Myscelia* TSDN spectral tuning, increasing by ∼15% to 0.72 in TSDN_H_, and ∼9% to 0.93 in TSDN_V_ (Fig. 6C, S9).

## Discussion

Overall, we observed a separation of spectral sensitivities in butterfly DNs that aligned with functional differences in motion sensitivity. Butterfly WFDN activity closely resembles those described in other species^43–55^, with a maximum sensitivity to wide-field motion along a preferred direction (Fig. 2&3). Butterfly WFDNs have effectively broadband spectral responses (Fig. 2-3), although subtle species-specific differences in WFDN spectral tuning corresponds with both the available set of photoreceptors (Fig. 1C) and the spectral sensitivity of optomotor head rotations (Fig. 5), supporting the involvement of WFDNs in controlling stabilisation reflexes^43^. In contrast to WFDNs, butterfly TSDNs are comparatively narrowband (Fig. 2&4) and match conspecific wing coloration (Fig. 6), suggesting a role in coordinating sexual pursuit behavior.

Indeed, butterfly TSDNs sample from the frontal visual field, aligning with optical acute zone specialisations for target tracking described in other Nymphalids^133, 134^. In sexually dimorphic butterflies, male wing coloration signals both sexual fitness to the female, and sexual identity to repel unwanted male advances^106, 107^. Whilst little is reported regarding *Myscelia* behavior, males in captivity readily engage in fast aerial pursuits and perch with their dorsal wing coloration displayed (Movie S3), similar to other species with prominent dorsal wing coloration^107^. The tuning of male *Myscelia* TSDNs to male wing coloration suggests a potential role in territorial pursuit, and future work should investigate whether female TSDNs have similar spectral selectivity. In *Danaus*, TSDNs in both sexes are exquisitely tuned to long-wavelengths (Fig. S4). Notably, wing coloration in *Danaus* is sexually monomorphic due to the constraint of Müllerian mimicry, whereby divergence from the population visual trait is penalised by increased predation^123, 125^. *Danaus* may therefore rely more on olfaction to determine mate suitability^135, 136^. Indeed, males engage in aerial ‘hair-pencilling’ to disseminate pheromones over the female antennae during courtship flight^98, 102, 104, 137^. In Noctuid moths, female pheromone produces upwind flight in males by amplifying WFDN sensitivity^52^ and optomotor reflexes^138, 139^; it would be interesting to investigate whether olfaction modulates butterfly TSDN responses in a similarly sexually dimorphic manner.

Butterfly TSDNs largely resemble those described in Diptera^40, 50, 63, 64^ and Odonata^58–61, 76^. However, whilst most TSDNs responded exclusively to moving targets, *Danaus* TSDN_D_ also responded to gratings moving in the preferred direction, albeit at a lower firing rate than typical of WFDNs. Butterfly TSDN size tuning is similar to that of Dipteran TSDNs^40, 50^, but are sensitive to larger targets than dragonfly TSDNs^76^. However, dragonfly TSDN target size preference ranges from 1°-diameter in bright skylight to 16° in a dim laboratory due to light adaptation^76^, thus butterfly TSDNs may display similar variability as a function of adapting background luminance. Notably, preferred target sizes were smaller for bright targets compared to dark targets (Fig. 4). The irradiation illusion is a similar phenomenon in human vision, whereby a bright object on a dark background appears larger than the same sized object with opposite contrast^140, 141^. This arises from a saturating non-linearity in ON-contrast sensitivity compared with near-linear OFF-contrast sensitivity^142, 143^. However, the magnification factor of the irradiation illusion in humans is ∼1.5x^141^, so other mechanisms may be involved in the ∼8x increase in preferred size observed in butterfly TSDNs.

Whilst butterfly TSDN spectral-, size-, and velocity-tuning suggests involvement in sexual pursuit behavior, flowers subtend similar visual angles as butterflies during flight, and it is likely that DN populations function collectively to support multiple behaviors to minimise energetic expenditure^50, 144^. Many insects display innate spectral preferences for various behaviors including feeding, mating, and oviposition^145–148^. Interestingly, whilst spectral preferences often differ between behaviors^108, 149^, some species, including *Danaus*, have an innate preference to feed from flowers spectrally similar to conspecific coloration^150, 151^. Innate spectral preferences readily shift to other wavelengths upon training^147, 151^, and coincides with spectral re-tuning of mushroom-body neurons^147^. Testing whether butterfly TSDNs display comparable spectral plasticity will provide insight into their involvement in controlling foraging behaviors.

Nonetheless, the difference in spectral sensitivities between butterfly WFDNs and TSDNs indicates a functional separation of spectral inputs to motion-vision in the context of stabilisation reflexes and target tracking, respectively. Diurnal insect optomotor systems primarily receive input from short visual fibre photoreceptors expressing broadband rhodopsin with sensitivity maxima for green^80–93^. This broad spectral sensitivity increases photon-catch, which in addition to the quadratic sensitivity to image contrast in elementary movement detection^152^, produces a stabilisation system robust to variations in image statistics whilst the animal moves through the environment^90^. Additional photoreceptors converge onto wide-field motion-vision in *Drosophila*^32^ and bees^77^, further improving the sensitivity and spectral range of optomotor reflexes^32^. Electrophysiological and connectomic data demonstrates that multiple photoreceptor inputs broaden the spectral sensitivity of most LMCs in *Papilio* butterflies^93, 122^. We found that *Myscelia* WFDN_H_ and optomotor responses could be well explained by predominantly green- and B^+^G^-^-photoreceptor input (Fig. S3). *Danaus* displayed a reduction in wide-field blue-sensitivity and an increased red-sensitivity that did not match green photoreceptor absorption (Fig. S3), and were best explained by dominant Y-receptor input, although other combinations produced similarly close fits of the data (Fig. S3D). Further analysis of WFDN_H_ and optomotor response half-saturation stimulus intensities^69, 85, 93, 145, 153, 154^, in combination with opsin photo-bleaching^155, 156^, will be required to unambiguously determine the photoreceptor identities contributing to wide-field responses in *Danaus*. Nonetheless, a broad spectral tuning of the optomotor system may be a general optimisation for producing consistent stabilising movements that are fast, sensitive, and robust to changes in the spectral content of the environment^90^.

However, what selective pressures drive the reduced blue-sensitivity and increased red-sensitivity of the wide-field system in *Danaus*? Possibly, the inclusion of motion-vision red-sensitivity comes at the cost of blue-sensitivity, or, alternatively, that the exclusion of blue-sensitivity increases red-sensitivity. Previous work in *Danaus* reported that (i) red-sensitive R9 inhibits G^+^R^-^-opponent R1-2, photoreceptors that are usually UV- or blue-sensitive in simple retinas, and (ii) that blue-sensitive photoreceptors are long-wavelength inhibited^118, 129^. Whilst long-wavelength vision in *Danaus* matches conspecific wing-coloration, red-receptors may have other general advantages for motion-vision. For example, foliage has broad reflectance extending towards long-wavelengths, with average flower reflectance continuing to increase beyond 700nm^157^. Further analysis of wide-field motion vision in butterflies with red-sensitive retinas will be required to determine the prevalence and functional advantages of motion-vision red-sensitivity, especially in species with non-red wing coloration such as *Papilio*^158^ or *Pieris*^159, 160^.

A previous investigation of TSDN spectral sensitivities reported adaptations to improve photon catch^69^, reminiscent of optomotor spectral-broadening. Dragonflies hunt from below to contrast flying prey silhouettes against the sky^72–74^, and their TSDNs are correspondingly aligned to sample the dorsal visual field^59–61^, and are spectrally tuned to UV/blue^69^. This matches an enrichment of UV/blue sensitivity in the dorsal retina^70^, along with other optical and physiological specialisations that enhance skylight photon-catch^75^. The narrowband spectral tuning of butterfly TSDNs suggests that, despite lower total photon-catch, target-tracking signal-to-noise is overall enhanced by band-pass spectral filtering matched to the coloration of interest. However, whether TSDN spectral-selectivity arises directly at the level of motion-detection in the optic lobe, or via convergence of chromatic input onto an otherwise achromatic motion-vision pathway remains to be tested. In *Sarcophaga* blowflies, male-specific TSDNs have secondary arborisations in the anterior optic tubercle^63^, a target neuropil for processing polarisation and chromatic outputs from the optic lobe^161, 162^, suggesting a potential convergence of parallel chromatic/achromatic inputs. However, asymmetric spectral-sensitivities of ON and OFF pathways in *Drosophila* motion-sensitive T4/T5 cells enhances the detection of approaching objects that differ in color to the background^20^, demonstrating advantages for intrinsically chromatic motion-vision. Notably, *Myscelia* TSDN responses to dark objects on bright backgrounds were less sensitive when the background was blue (Fig. 4Aii), opposite to the maximum sensitivity to bright blue objects moving on a dark background (Fig. 4Aiv). This could indicate differences in ON/OFF spectral sensitivities in the target-tracking pathway, analogous to that described in *Drosophila* T4/T5 cells^20^. Future work should focus on determining the origin of spectral-selectivity in butterfly TSDNs, and modelling target-tracking performance in both direct and convergent configurations. This will provide potentially generalisable insights into the advantages of integrated color- and motion-vision, with applications to artificial vision systems.

## Methods

### Animals

Monarch (*Danaus plexippus*; Subfamily: *Danainae*, Tribe: *Danaini*) and Blue-Wave (*Myscelia cyaniris*; Subfamily: *Biblidinae*, Tribe: *Epicaliini*) butterflies were purchased as pupae from Stratford Butterfly Farm (UK) and The Entomologist Ltd. (UK). Pupae were stored at 23°C and 60% humidity on a 12-hr dark/light cycle. Imagoes were allowed to harden for 24-hrs post-emergence, then stored at 15°C. Butterflies were fed diluted honey every other day, and all experiments were performed within two weeks post-emergence.

### Wing Reflectance Measurements

Wings were amputated and flattened between two microscopy slides. The wing was illuminated by a Xenon short-arc bulb (SLS205, Thorlabs, USA) via a 200µm fibre and lens shutter which produced a ∼2mm diameter collimated light beam. Light was incident at an angle of 40° and measured using a calibrated Flame USB spectrometer (Ocean Optics, USA). Throughout this paper, ‘reflectance spectrum’ refers to the fraction of illuminant photon radiance reflected as a function of wavelength, and is assumed to be independent of the illuminant spectrum at the tested range of intensities. A ‘reflection spectrum’, in contrast, is defined as the spectrum of photon radiance reflected from the wing, which does depend on the illuminant spectrum. Under white light illumination, the normalised reflection spectrum corresponds to the normalised reflectance. Under any other illuminant, the reflection spectra is a function (assumed here as multiplication) of the reflectance and the illuminant spectrum. Natural reflection spectra were estimated by multiplying the standard wing reflectance with the CIED65 standard daylight illuminant spectrum (ISO/CIE 11664-2:2022; International Commission on Illumination, Commission Internationale de l’Eclairage, CIE).

### Visual Stimuli

For descending neuron electrophysiological experiments, butterflies were positioned with their head centred 12 cm from a 16 x 9 cm (width x height) rear projection screen (ST-Pro-X, Screen-Tech e.K., Hohenaspe, Germany), thereby subtending 67° x 41° (azimuth x elevation) of the frontal visual field. A customised display system comprising a multi-LED galvo scanner and an RGB DLP projector was focussed onto the rear surface of the projection screen (Fig. S1).

The multi-LED galvo scanner AT20L-L (Shenzhen City Aoxinjie Technology, China) was illuminated by six LEDs (LED-Engin, ams-OSRAM, Austria; peak wavelengths: 395/450/499/518/595/636nm; Fig. S9) transmitted via a 5 mm diameter liquid light guide (Newport, USA). The light guide output was focussed onto the rear surface of the projection screen via an EF 100mm f/2.8L Macro IS USM lens (Canon, Japan), generating an 8°-diameter circular bright target on a dark background. The horizontal and vertical position of the target was controlled by two orthogonal galvo-mounted mirrors (RGB-Lasers, Netherlands). Each galvo motor was controlled by a driver with ±10 V input. Voltages were generated in Spike2 software and output from a Micro 1401 ADC (CED, UK) at 1 kHz temporal resolution. Linear voltage ramps moved the bright target across the screen at 115 °/s. For each trajectory, the target first remained stationary at the start position for 500 ms to exclude transient responses to target appearance from the measurement. The color of the moving target was varied sequentially in ABBA sequence to mitigate time-dependent changes in the adaptation state of the neuron. LED emission intensities were measured using a calibrated spectrometer (Flame USB, Ocean Optics, USA). Linear regressions of the integrated LED spectra photon count vs input pulse-width modulation (PWM) were calculated and used to equalise the output of each LED to match the maximum output of the dimmest LED.

The RGB DLP projector (Lightcrafter 3010, Texas Instruments, USA) was used to present different sequences of moving targets (including variable size/velocity) and gratings to discern cell types. Moving patterns were generated at 1280 x 720-pixel resolution in monochromatic mode with 8-bit depth to increase the native DLP framerate to 360 Hz. The intensities of each RGB LED were equalised by adjusting the supply current using the same calibration procedure as used for the LED-Engin (Fig. S1). DLP Local Area Brightness Boost was enabled and set to strength and sharpness of 0, and Content Adaptive Illumination Control was disabled to ensure that the calibrated LED intensities were used. Moving patterns were generated in MATLAB using PsychToolBox^163^, rendering sequential patterns onto each RGB subframe to achieve 360 Hz frame rates^164^. Visual stimuli timings were aligned with electrophysiology via a photodiode measuring a small 100-pixel square in the corner of the screen which alternated on/off every frame.

Descending neuron cell types were classified using a sequence of standardised green visual stimuli presented in the frontal visual field using the DLP projector, comprising: (i) wide-field gratings moving sequentially in eight directions (10° spatial wavelength, 5 Hz temporal frequency, unless specified otherwise); (ii) rasterised scans of a small circular (4° diameter) target moving sequentially along eight directions to determine the spatial receptive field and preferred direction; (iii) square targets of varying size moving through the neuronal receptive field in the preferred direction. For size tuning, the targets moved at a constant velocity of 115°/s. For velocity tuning, the square target side lengths subtended 4°. Both target size and velocity tuning experiments presented each variable in ABBA sequence to mitigate time-dependent changes in the adaptation state of the neuron. For all moving target stimuli, the target appeared stationary at the start position for 150 ms prior to movement to exclude transient neuronal responses to target appearance. For color experiments, gratings were always either blue, green, or red bright bars presented on a black background.

For optomotor experiments, butterflies were positioned with their head centred 10 cm from a 54 x 30 cm, 1920 x 1080 pixels (width x height) 240 Hz LCD computer monitor (S25BG400EU, Samsung, South Korea), thereby subtending 139° x 113° (azimuth x elevation) of the frontal visual field. Visual stimuli were generated at 240 Hz in MATLAB using PsychToolBox^163^. The intensity of each RGB color channel was equalised by adjusting the greyscale values, using the same linear regression calibration protocol as described for the LED-Engin. Moving gratings were presented sequentially for 2 seconds in both left and right directions with 20° spatial wavelength and temporal frequencies ranging from 0.1 Hz to 40 Hz. A small square trackbox in the corner of the screen turned white when stimuli were running to align the stimuli with stereo camera footage. An IR photodetector (Stock # 642-4430, RS Components Ltd, UK) measured changes in trackbox intensity via an Arduino and was used to toggle an IR LED on/off within the stereo camera field of view.

### Photoreceptor Electrophysiology

Spectral sensitivity of the photoreceptors was measured with single-cell recordings of membrane potential during stimulation with iso-quantised flashes, produced with a Xe arc lamp (Cairn, UK) and a monochromator (B&M Optik, Germany) or with a multispectral LED array^165^. The method is described in detail elsewhere^118, 129^. Photoreceptor identity was determined via angular maximum of polarization sensitivity (vertical for R1,2,9; horizontal for R3,4; diagonal for R5-8). For both species, animals of both sexes were surveyed. Each electrode excursion yielded many tens of impaled photoreceptors, which were quickly scanned with LED array. Finally, min. 4 cells per spectral class were scanned with the monochromator and included into the spectral sensitivity graph.

### Descending Neuron Electrophysiology

Butterflies were mounted onto a large rotatable platform to position the screen in the frontal visual field of the butterfly (defined as parallel to the flat caudal surface of the butterfly head capsule). The ventral nerve cord was accessed ventrally, and a hypodermic needle fashioned into a hook supported the nerve during electrophysiological recordings. 3-MOhm sharp tungsten electrodes (UEWSHGSE3P1M, FHC Ltd, USA) were used to measure extracellular descending neuron action potentials, which were amplified at 10k gain with an extracellular amplifier (DP-314, Warner Instruments, USA) and digitised at 25 kHz using a Micro 1401 (CED, UK), and spike-sorted in Spike2 (CED, UK; Fig. S2). Spike sorting quality was assessed from inter-spike interval histograms (with events less than 2ms classified as false positives), and time-continuous principal components of waveform shape (Fig. S2).

### Modelling DN Responses from Photoreceptor Inputs

Photoreceptor responses were modelled following previous protocols^157, 166, 167^. Normalised photoreceptor voltage, *V*, was given by the log-logistic relation:

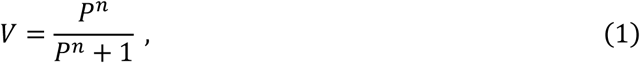

where *P* (photon-catch) is the relative amount of light absorbed by each photoreceptor (normalised to the adapting background light level, below), and n is the Hill coefficient. Equation (1) therefore resembles the characteristic photoreceptor Voltage-log_10_(Intensity) contrast-response function^167^. A Hill coefficient of 1 was used for all photoreceptors, corresponding to a dynamic range of ∼2 log units. Normalised photon-catch, *P*, was calculated as:

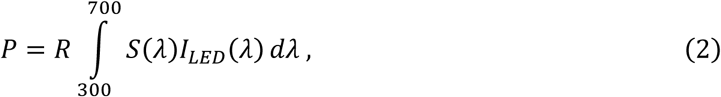

where *λ* is wavelength in nm, *S*(*λ*) is the normalised photoreceptor spectral sensitivity, and *I_LED_*(*λ*) is the LED emission spectra (photons/cm^2^/s). The range-sensitivity, *R*, scales the photon-catch dynamic range, and is modelled as the reciprocal photoreceptor photon-catch at the adapting background intensity, I_BG_ (photons/cm^2^/s):

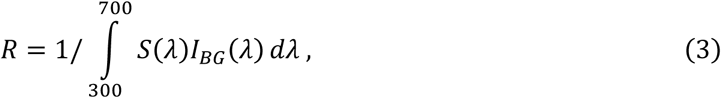

where *R* is formulated such that the adapting background photon radiance produces a normalised photoreceptor photon-catch of P = 1 from equation (2), corresponding to a photoreceptor half-max response V = 0.5 from equation (1), in accordance with the observation that photoreceptors maximise their information capacity by centring their dynamic range on the adapting background intensity^167, 168^.

Photoreceptor responses were calculated for each of the six LEDs used for DN spectral tuning experiments. Best-fit photoreceptor weights were obtained by calculating a least-squares linear regression of the measured DN responses as a function of photoreceptor activation to the same LEDs, i.e.:

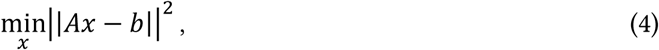

where A is the matrix of photoreceptor responses to each LED (with dimensions: #LEDs x #photoreceptors), x is the vector of photoreceptor weights, and h is the vector of DN responses to each LED. Solution stability was assessed by comparing regularised fits with variable penalties for photoreceptor weight magnitudes, i.e.:

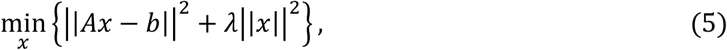

where A is the regularisation penalty weight.

### Optomotor Head Rotations

Butterflies were tethered in front of a computer monitor in a dark arena by supergluing a 1 mm diameter steel rod to the mesoscutum and were encouraged to hold onto a small ∼3x3 cm piece of tissue paper as a perch throughout the experiment. Head rotation was measured using two frame-synchronised FLIR Blackfly S USB3 0.4 Megapixel cameras (Teledyne, Canada) recording at 250 frames per second using SpinView (Teledyne, Canada). Cameras were fitted with 6 mm fixed focal length lenses (UC Series, Edmund Optics, USA) with aperture set to f/5.6. To reduce luminance fluctuations from the visual stimulation, both lenses were fitted with an IR long-pass filter (25mm diameter, Stock #12-766, Edmund Optics, USA), and the butterfly was illuminated with IR LEDs (Model #S1120B, Splenssy). For *Danaus*, three natural circular head markings on the dorsal surface of the head capsule (vertex) were used to measure head orientation as they form a plane parallel to the transverse plane of the eye. In *Myscelia*, where natural head markings are absent, a small strip of retroreflective tape (ualisys, Sweden) was mounted to the head to form the left-right lateral axis, and a small spot of white nail polish marked the rostral-caudal axis. Head markings were tracked using DeepLabCut^169^. Stereo-cameras were calibrated using a 7 x 6 checkerboard (side length = 3.126 mm), and stereo pairs of 2D xy pixel coordinates were triangulated into 3D xyz coordinates using Anipose^170^. Head orientation was calculated relative to the neutral head orientation, defined as the average across each ∼5-minute recording. For each time point, the relative rotation matrix to align the local head coordinate system with the neutral head reference coordinate system was calculated using the dot product via the yaw-pitch-roll Tait–Bryan sequence. Head yaw-pitch-roll angles were subsequently smoothed using a Savitzky-Golay filter (3^rd^ order, 404 ms window).

### Data Analysis and Statistics

All data analysis was performed in MATLAB (The MathWorks, USA). For all DN tuning curves, responses were calculated as the average spike rate during pattern movement. DN receptive fields were calculated by extracting the position, and direction of target movement at the time of each spike. For each of the eight directions of target motion, pixels corresponding to the spike-triggered target location were transformed into visual azimuth/elevation, binned into a 5°x5° grid, and smoothed using a gaussian kernel (1 standard deviation). Local motion sensitivity and preferred directions were calculated for each bin using the least-square sinusoidal fit of responses to each target direction.

WFDN_H_ spectral tuning curves were analysed using a one-way ANOVA, making the conservative assumption of independence between the six LED groups. Spectral tuning curves were normalised to the mean across LED conditions to remove variation in absolute spike rates between individuals. Statistically significant differences in the ANOVA results were further analysed by post-hoc t-test comparisons to the average response across LEDs.

Due to violations of normality and equality of variance, statistical analyses of DN tuning curves for gratings direction, target size and velocity were performed using non-parametric permutation tests comparing the observed pairwise mean differences between color conditions to the empirical null distribution of pairwise differences, H_0_^171, 172^ (Fig. S5-5). For each animal, spike rate tuning curves were normalised by the min-max across color conditions. To generate H_0_, pairwise differences were calculated from data pooled from the three color conditions across N animals, yielding (3!)^N^ = 6^N^ unique pairwise mean differences. The probability of the observed mean differences occurring under H_0_ (i.e., p-value) corresponds to the fraction of mean differences that are more extreme than the observed difference. To control for false positives from multiple testing across the independent variable range (i.e. gratings direction, target size, or target velocity, for which responses are correlated), H_0_ was calculated separately and z-standardised for each variable, and the largest z-value across the independent variable for each permutation was stored to generate a max-value distribution. The 95^th^ percentile of this max-value distribution was used as an adjusted significance threshold, and corrected p-values were calculated as the fraction of max-values more extreme than the observed mean differences (Fig. S5-5). Finally, false positives from multiple testing across color conditions was controlled by applying the Bonferroni correction (three tests).

Optomotor tuning curves were mean normalised across color conditions for each animal, then normalised to the maximum average response across color conditions (green in both species). Individual responses at the preferred temporal frequency of the population average across color conditions (20 Hz in both species) were analysed using a repeated-measures ANOVA of color conditions, followed by pairwise t-test comparisons with Bonferroni correction. Non-normalised raw data was non-normally distributed (Lilliefors goodness-of-fit, p>0.05), and therefore analysed using a repeated-measures Friedman test (Fig. S7).

Head movements to moving objects were normalised to the head orientation at the start of object movement. The maximum change in head yaw orientation towards the direction of target motion was pooled from all trials (Fig. S8C). The 95^th^ percentile of the cumulative distribution function (CDF) of maximum yaw values was used as a threshold to identify trials in which the animal tracked the object. Pair-wise differences between CDFs of yaw responses to different colors were analysed using two-sample Kolmogorov-Smirnov tests for each tested object size. P-values were adjusted with Bonferroni correction for color conditions.

Spectral alignment between DN spectral tuning curves and (i) wing reflectance/reflections (Fig. 6), and (ii) modelled photoreceptor combinations (Fig. S3) was calculated as the Pearson correlation coefficient (i.e. cosine similarity) between 395-636nm (i.e., the range used to measure DN spectral tuning). For wing reflectance/reflection comparisons, TSDN spectral tuning curves were linearly interpolated and smoothed (Savitzky-Golay filter, 3^rd^ order) with filter windows varying from 11 to 241 nm (Fig. S9).

## Supporting information

Movie_S1_Dplexippus_optomotorResponse

Movie_S2_Mcyaniris_optomotorResponse

Movie_S3_Mcyaniris_slow5x

## Acknowledgements

This work was supported by Air Force Office of Scientific Research (AFOSR) grant FA8655-23-1-7049 to GB and HGK, Biotechnology and Biological Sciences Research Council (BBSRC) grant BB/X002276/1 to HGK, and Bioinspired Sensing Computing and Control with International Teams (BISCCITs) travel grant to JAS. Thanks to Ric Wehling and Nandini Iyer for providing financial support.

---

**Figure S1:**
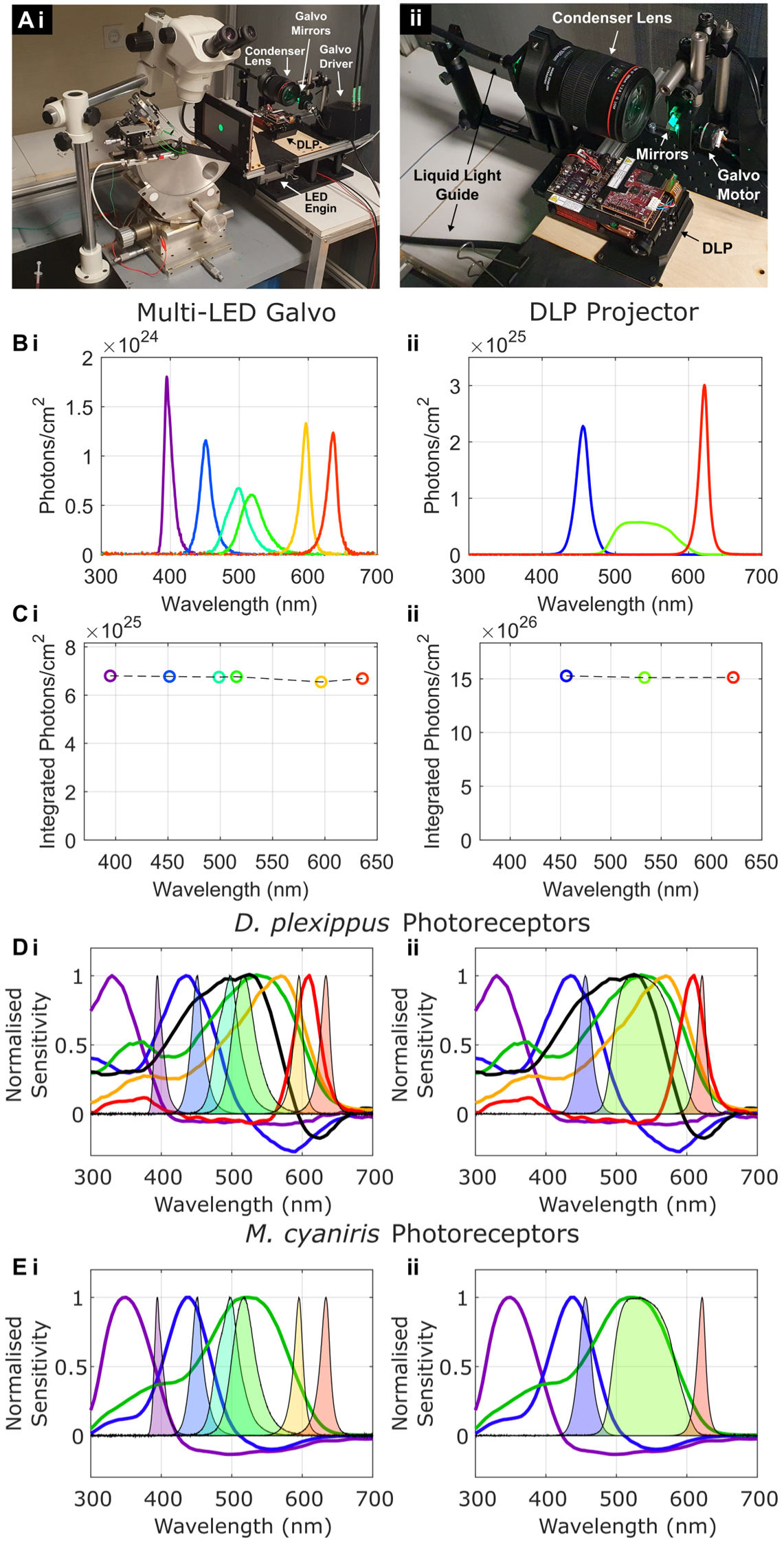
Multi-LED Galvo display system and LED calibration (Related to Figure 2) (A) Photographs of the electrophysiology set-up. (i) The butterfly is mounted on a rotation platform with the projection screen positioned in the frontal visual field. An example green target subtending 8°-diameter on the butterfly retina is presented from the multi-LED galvo system. (ii) Details of the display system. The galvo XY mirrors are in line with the DLP projection axis. A liquid light guide directs light from the LED Engin (out of view) into a Canon 100mm Macro lens acting as a condenser. (B) Calibrated LED spectra for the (i) multi-LED galvo system, and (ii) the DLP projector. (C) Integrated photon intensities for each calibrated LED. (D) Comparison of normalised LED spectra with *D. plexippus* photoreceptors. (E) Comparison of normalised LED spectra with *M. cyaniris* photoreceptors.

**Figure S2:**
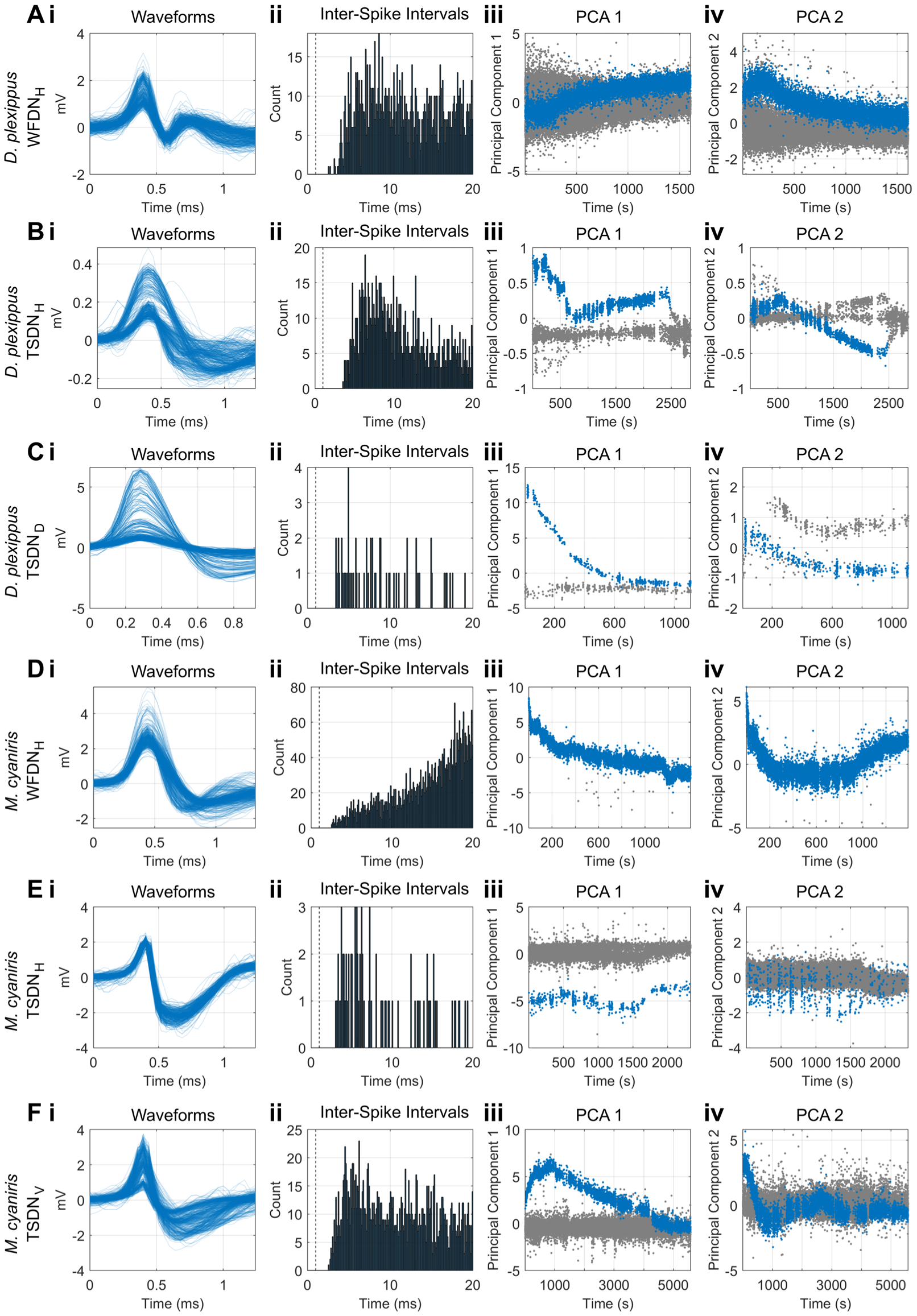
Extracellular electrophysiology and spike sorting (Relating to Figures 2-4) Example spike-sorted DNs from **(A-C)** *D. plexippus*, and **(D-F)** *M. cyaniris*. (i) Random subset of 500 waveforms. (ii) Inter-spike interval histograms. Dashed line at 1ms. (iii) First principal component (PC1) of waveform shapes vs recording time. (iv) Second principal component (PC2) of waveform shapes vs recording time. Spike waveform shapes often drifted throughout recordings due to changes in electrode positioning and ventral nerve cord hydration, however the same isolated unit could be identified due to the time-continuity of this drift in PC 1-2 space.

**Figure S3:**
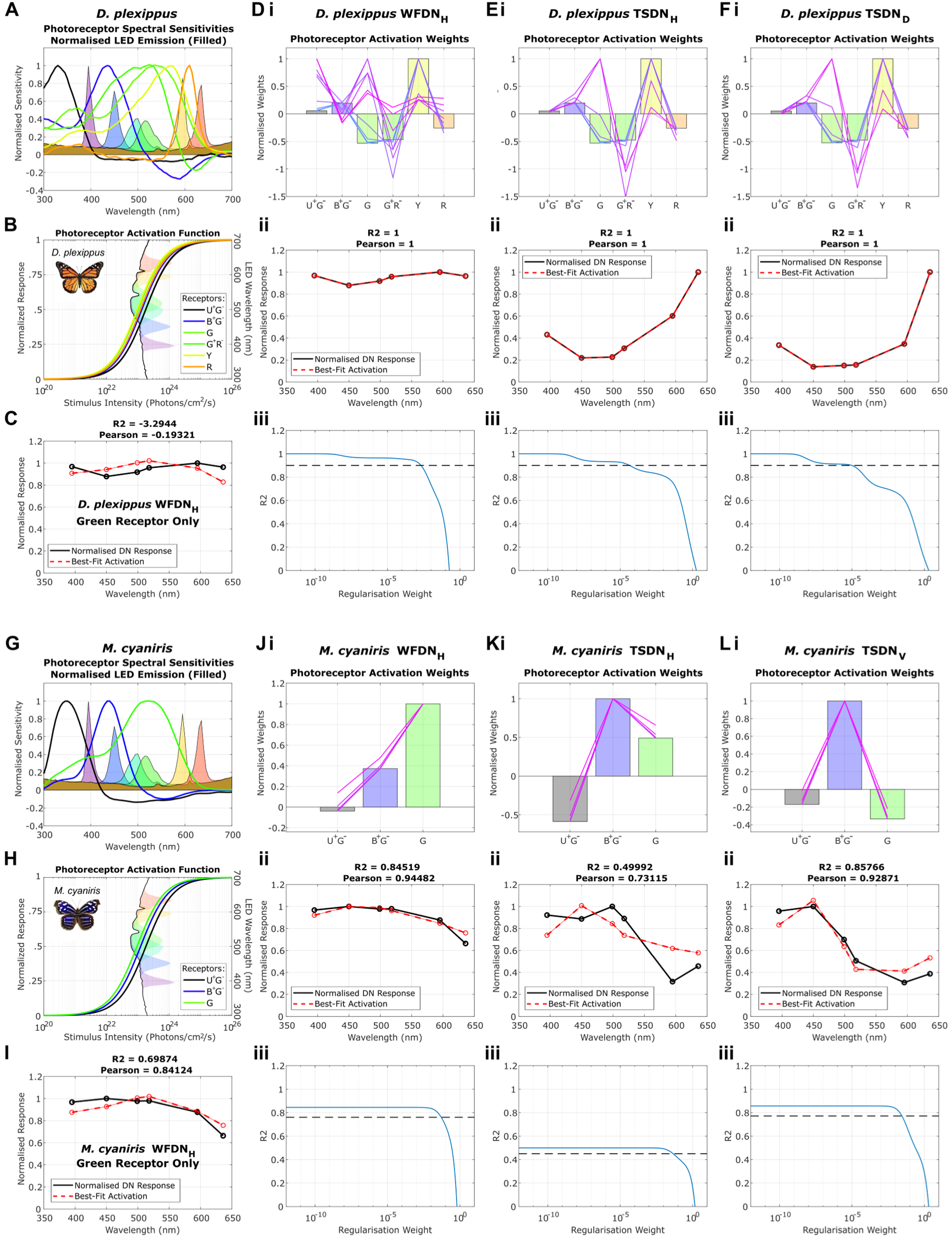
Modelling DN responses from linear combinations of photoreceptor inputs (Relating to Figures 1-2). (A) Normalised photoreceptor spectral sensitivities (reproduced from Figure 1) and normalised isoquantal LED emission spectra (filled curves) used for the multi-LED galvo stimulus. The filled brown area indicates the background photon radiance to which photoreceptors are assumed to be adapted. (B) Log-logistic normalised photoreceptor activation as a function of light intensity, i.e. photons/cm^2^/s. Absolute LED emission intensities are plotted against wavelength on the right Y-axis. Black line corresponds to the background intensity (photons/cm^2^/s) to which photoreceptors are assumed to be adapted. (C) Least-squares best-fit WFDN_H_ response (dashed red) from green receptor input only, compared with the measured normalised WFDN_H_ response (black; reproduced from Figure 2). (D) Least-squares best-fit of WFDN response using a linear combination of photoreceptor inputs. Un-regularised and regularised photoreceptor weights are compared to indicate the stability of the simulation. Larger variability of regularised weight combinations indicates ill-conditioned solution instability with high sensitivity to measurement noise. (i) Bars correspond to the best-fit un-regularised relative photoreceptor weights (activations). Overlayed lines indicate the best-fit regularised photoreceptor weights (minimised weight magnitudes-squared) that produce simulated DN responses with R^2^ ≥ 90% of the un-regularized R^2^ value. Regularised weights are color-coded blue-to-magenta corresponding to progressively increasing penalties on photoreceptor weight magnitudes (and consequently have progressively decreasing R^2^ values). (ii) Least-squares best-fit WFDN_H_ response (dashed red) from the unregularized photoreceptor weights in (i), compared with the measured normalised WFDN_H_ response (black; reproduced from Figure 2). R^2^ and Pearson Coefficients indicated. (iii) Variation of simulated DN fit R^2^ values as a function of regularisation weight. Larger regularisation weight corresponds to a higher penalty on photoreceptor weight magnitudes. Dashed black line indicates the 90% maximum R^2^ value. The corresponding regularised photoreceptor weights producing R^2^ values above this threshold are plotted in (i). (E) *D. plexippus* TSDN_H_. (F) *D. plexippus* TSDN_V_. (G-L) Same analysis for *M. cyaniris.* Note that regularised solutions are less variable for *M. cyaniris* compared with *D. plexippus*, indicating solution stability.

**Figure S4:**
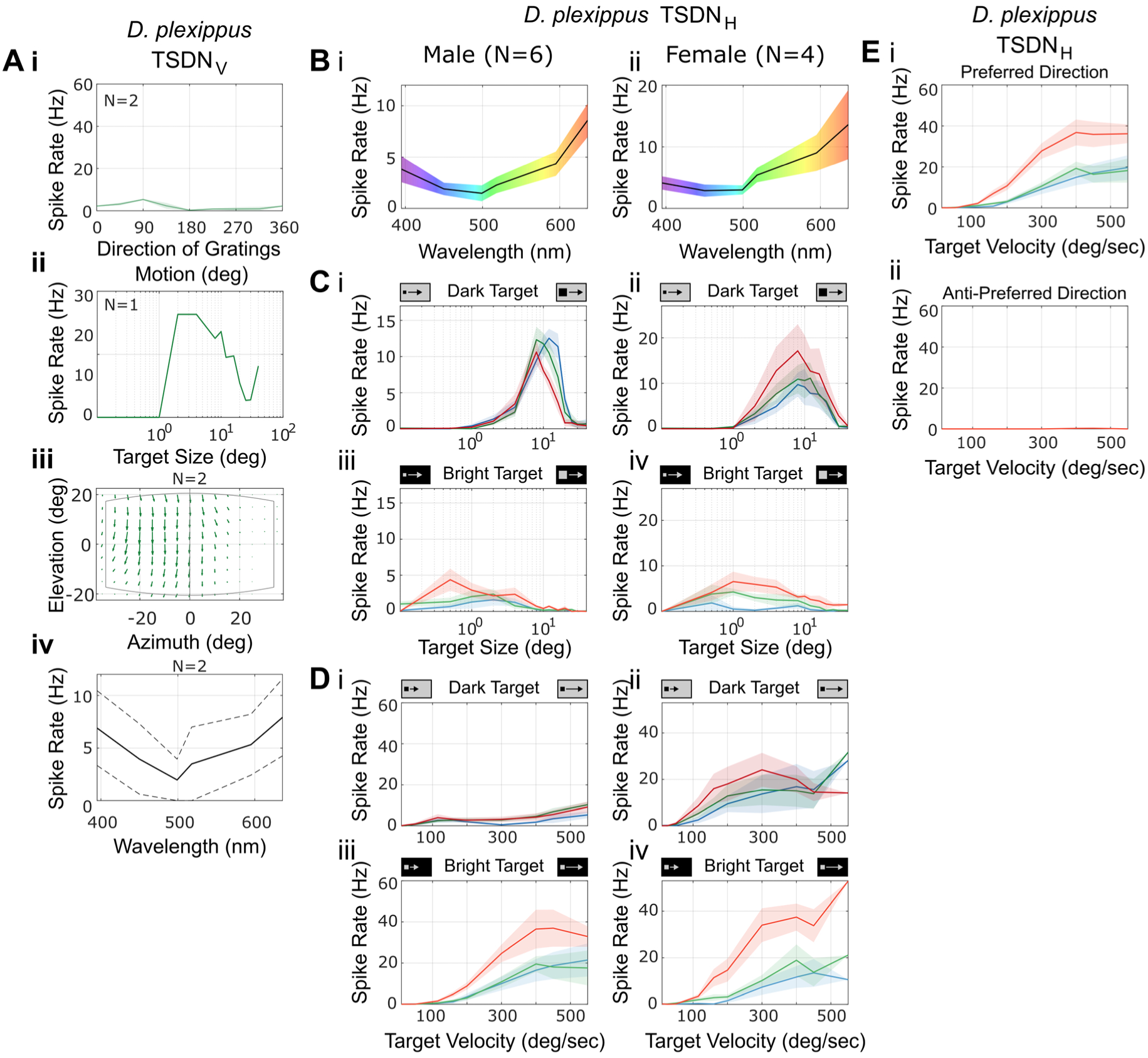
*D. plexippus* TSDN_V_, Comparison of *D. plexippus* TSDN_H_ Sex-Specific Responses, and Verification of directional selectivity at high velocities (Relating to Figures 2&4) **(A)** *D. plexippus* TSDN_V_ responses to (i) moving gratings (n=2), (ii) square dark targets of different sizes moving in the preferred direction (n=1), (iii) receptive field (n=2), and (ii) spectral tuning (n=2, individuals in dashed black, average in solid black). Stimulus parameters the same as Figure 2. **(B-D)** Comparison of *D. plexippus* TSDN_H_ properties in (i) male (N=6 animals) and (ii) female (N=4 animals). (B) spectral tuning, (C) target size tuning, (D) target velocity tuning. Same data is plotted as combined averages in Figure 2 and 4. (E) *D. plexippus* TSDN_H_ responses to bright square 4° (side-length) targets moving in the (i) preferred and (ii) anti-preferred direction (N=5 cells from 4 animals). Panel (i) is reproduction of Figure 4Biii for comparison. Only responses to bright red objects shown in (ii). All plots show mean ± SEM.

**Figure S5:**
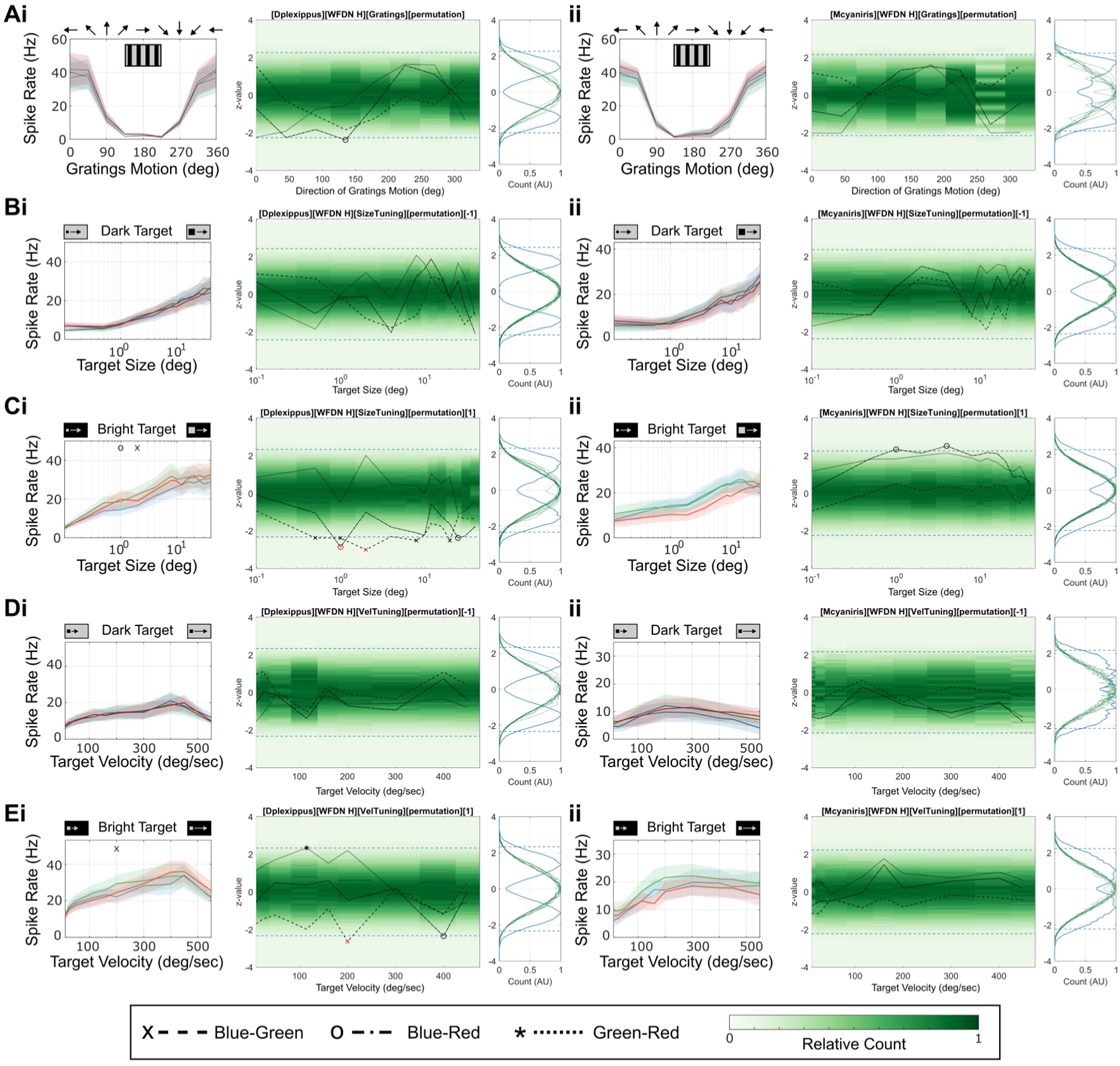
Non-parametric permutation analysis of Butterfly WFDN responses (relating to Figure 3) (Left) Tuning curves (reproduced from Figure 3) and (middle-right) permutation tests for (i) *D. plexippus*, and (ii) *M. cyaniris* WFDN_H_ responding to (A) moving gratings; (B) dark targets and (C) bright targets of different size; (D) dark targets and (E) bright targets of different velocities. Note that only velocities ≤450°/sec were included in permutation analyses as not all animals were tested with higher velocities. See Figure 3 for experiment details. Middle panel shows the z-standardised empirical null distribution H_0_ (green) for each independent variable value (i.e. gratings direction, target size, or target velocity). Right panel shows the same z-standardised H_0_ distributions overlaid (green lines). The symmetric distribution in blue shows the histogram of maximum absolute difference values that occurred in each permutation. The significance threshold (dashed blue lines) controls for the family-wise error rate across the (correlated) independent variable, and corresponds to the 95^th^-percentile of the blue max-distribution. The observed pairwise firing rate differences between the three color conditions are plotted over H_0_ in the middle panel (dash: Blue-Green; dot-dash: Blue-Red; dots: Green-Red). Black symbols (‘x’: blue-green; ‘o’: blue-red; ‘*’: green-red) plotted beyond the max-value threshold (dashed blue) indicate observations that have a p≤0.05 of occurring were there no difference between responses to different colors, after correcting for multiple comparisons across the correlated independent variable. Red symbols indicate data that remains p≤0.05 after Bonferroni correction for multiple pair-wise comparisons across color conditions and correspond to the symbols in the tuning curves (left panel, data reproduced from main Figure 3). See methods for additional details.

**Figure S6:**
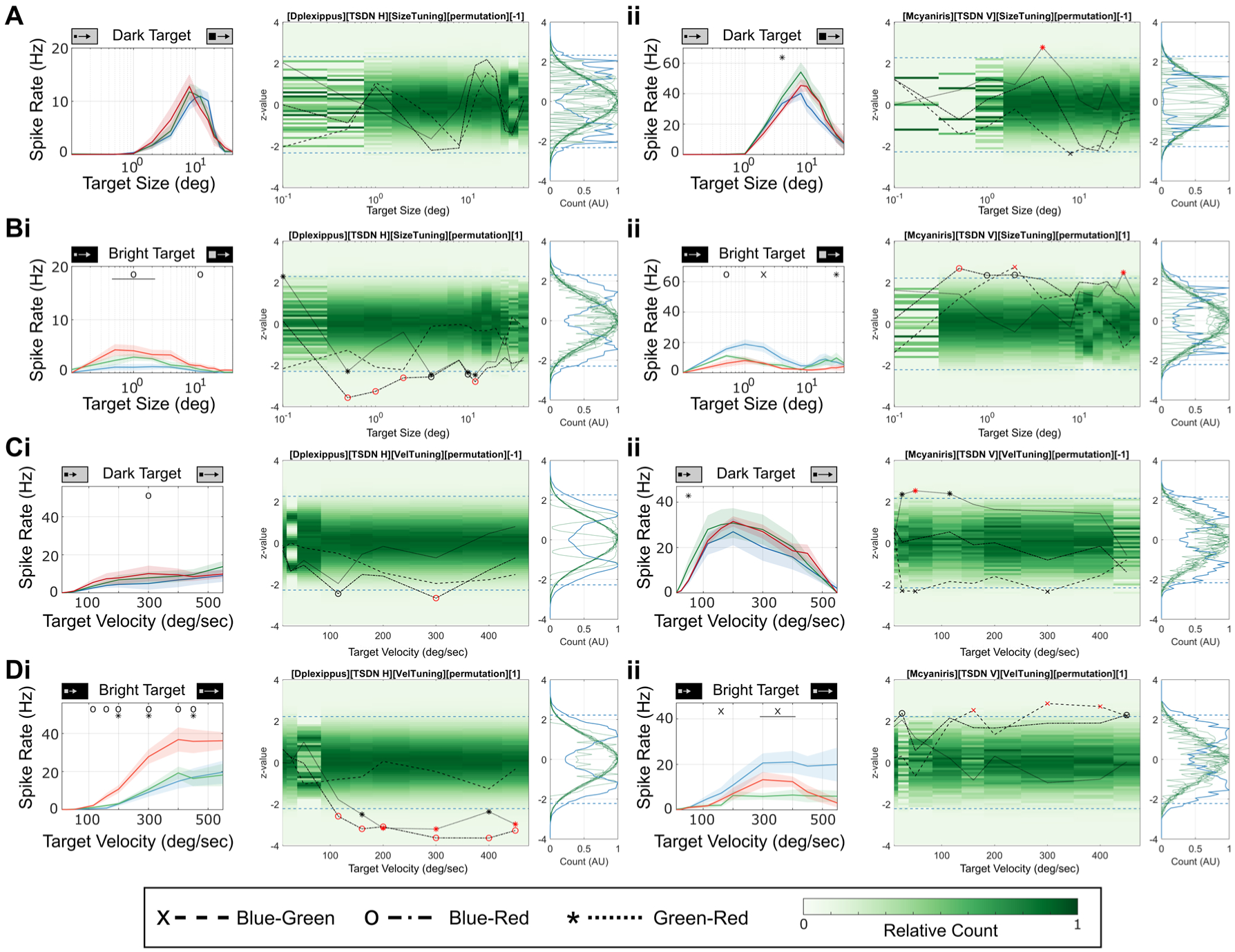
Non-parametric permutation analysis of Butterfly TSDN responses (relating to Figure 4) (Left) Tuning curves (reproduced from Figure 4) and (middle-right) permutation tests for (i) *D. plexippus* TSDN_H_, and (ii) *M. cyaniris* TSDN_V_ responding to (A) dark targets and (B) bright targets of different size, and (C) dark targets and (D) bright targets of different velocities. Note that only velocities ≤450°/sec were included in permutation analyses as not all animals were tested with higher velocities. See Figure 3 for experiment details. See Figure S5 for panel descriptions.

**Figure S7:**
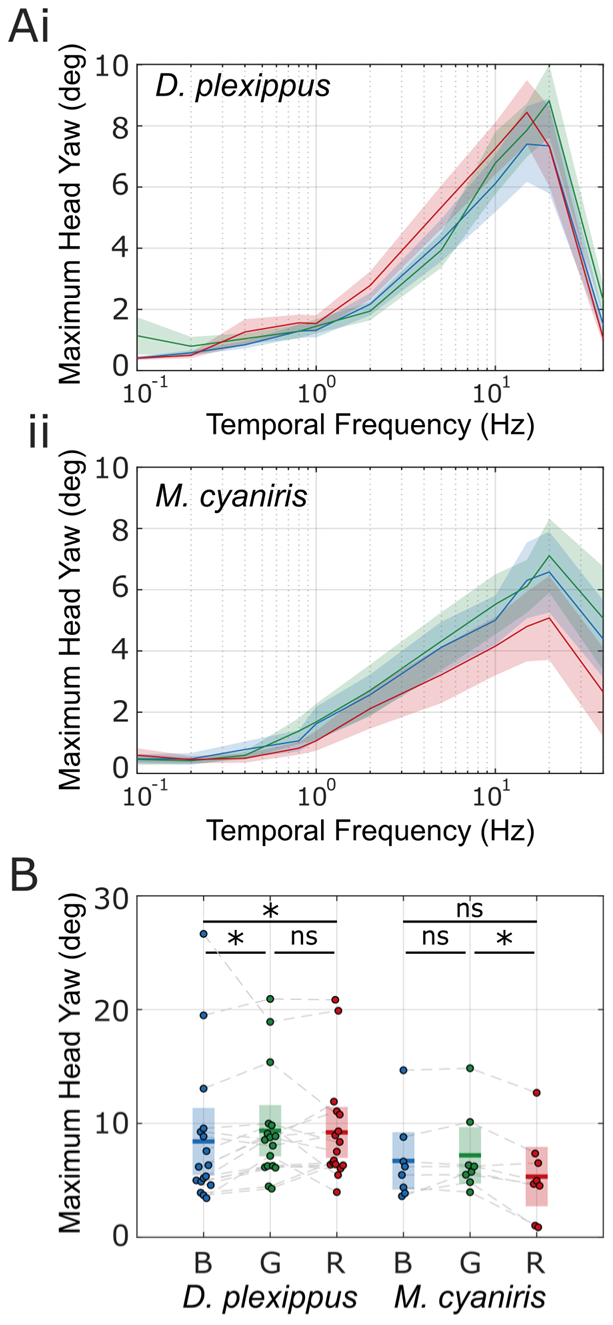
Raw data and statistical analysis of Butterfly optomotor head movements (Relating to Figure 5) (A) Raw maximum of averaged left and right head yaw amplitudes for each tested temporal frequency for (i) *D. plexippus* (N = 17 animals) and (ii) *M. cyaniris* (N = 8 animals). (B) Comparison of normalised maximum head yaw amplitudes for each grating color. Peak head yaw responses for different grating colors were statistically significantly different for both *D. plexippus* and *M. cyaniris* (repeated-measures Friedman test, p=0.011 and 0.044, respectively). Post-hoc Bonferroni-corrected pairwise comparisons indicated as * for p<0.05, and ns for p>0.05. *D. plexippus*: blue-green: p=0.049; blue-red: p=0.018; green-red: p>0.5. *M. cyaniris*: blue-green: p>0.5; blue-red: p>0.5; green-red: p=0.037.

**Figure S8:**
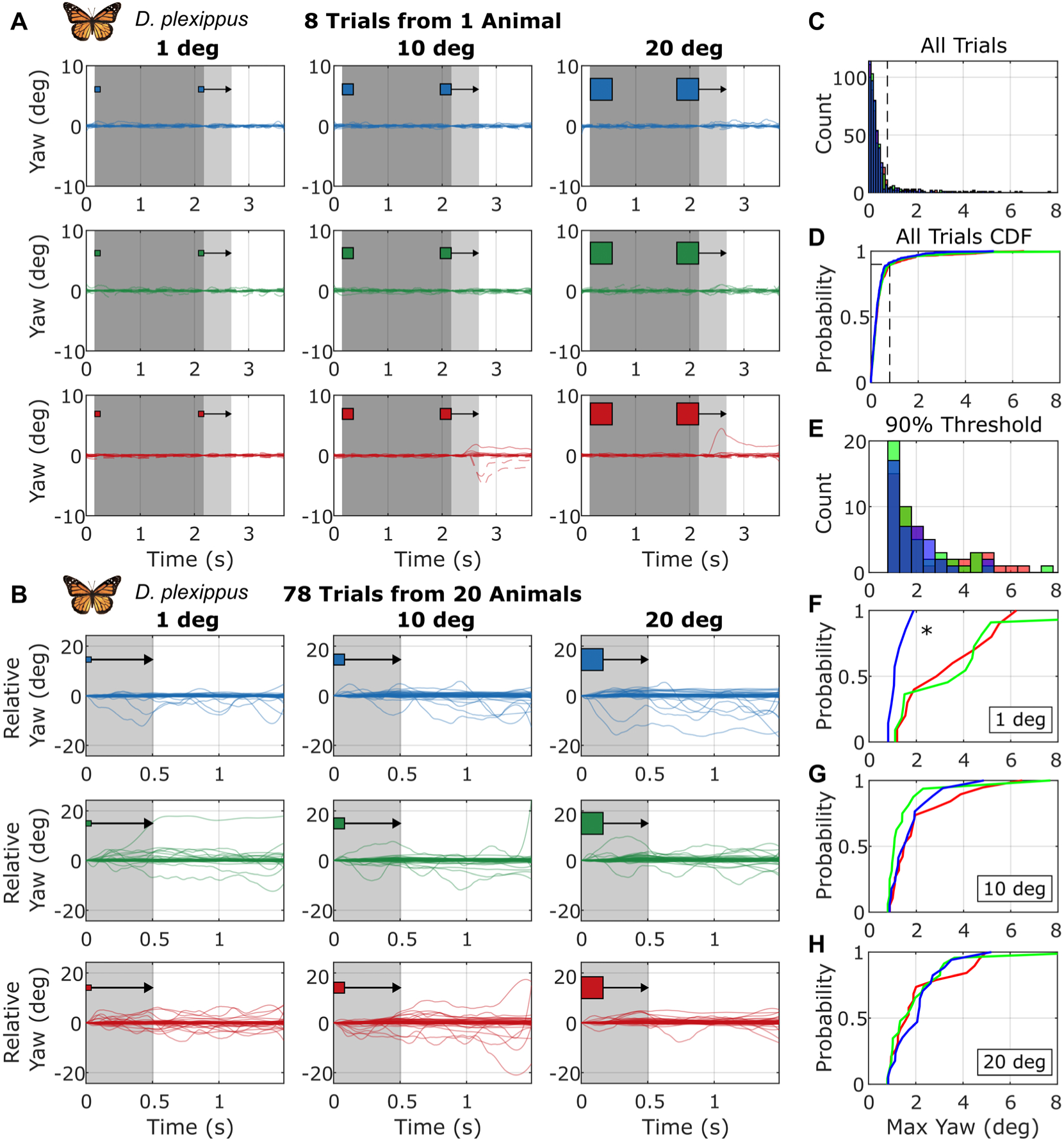
Head movements to moving targets in *D. plexippus* (Relating to Figure 5) **(A)** Example *D. plexippus* head yaw movements to horizontally moving targets of varying size (columns) and color (rows top/middle/bottom = blue/green/red). In each trial, a square target appears at one side of the screen and remains stationary for 2 seconds (dark grey area). The object then moves horizontally at 400°/sec (light grey area). Solid and dashed colored lines indicate responses to targets moving left and right, respectively. **(B)** Combined trials from 20 animals, only showing periods of target movement (light grey area). Head angles are normalised to the position at the onset of target movement. Responses to leftward targets are flipped, so all movements towards the direction of object motion are positive. **(C)** Histogram of maximum relative yaw angles (degrees) during the period of target movement in (B). Note subplots C-H share the same x-axis. **(D)** Cumulative Distribution Function (CDF) of maximum yaw angles for all blue/green/red target trials. The 90^th^ percentile is calculated and used as a threshold in plots (E-F). **(E)** Histogram of maximum relative yaw angles (degrees), excluding trials with amplitudes less that the 90^th^ percentile threshold. **(F-H)** CDFs of thresholded yaw amplitudes for blue/green/red targets with diameters of (F) 1°, (G) 10°, and (H) 20°. Asterisk indicates statistically significant differences between blue-green (pairwise Kolmogorov– Smirnov test, p=0.034) and blue-red (p=0.017) 1° diameter targets, without correcting for multiple comparisons. Differences were no longer statistically significant (p>0.05) after Bonferroni correction.

**Figure S9:**
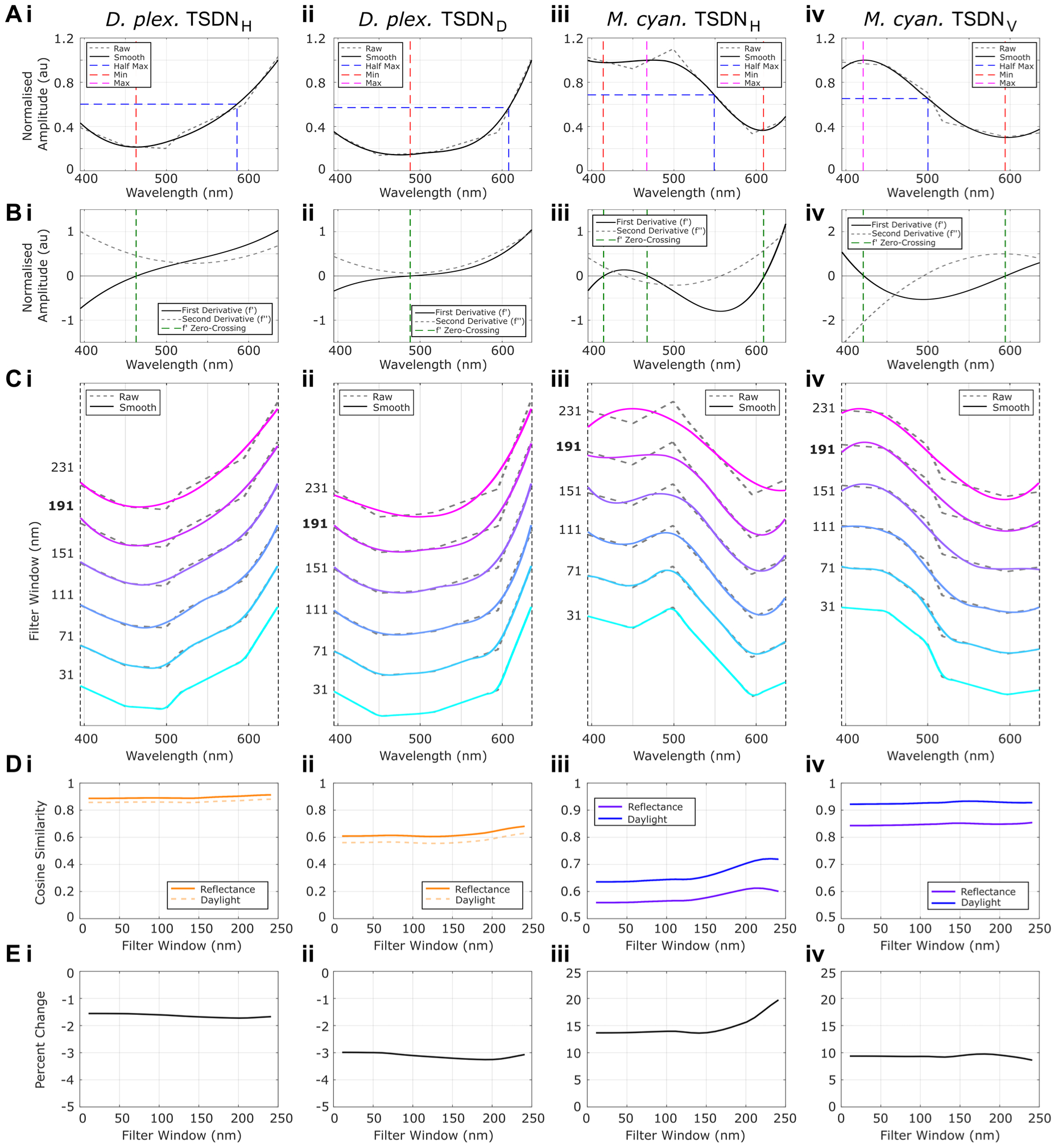
TSDN Parameterisation, Spectral Tuning Smoothing, and impact on Wing Reflectance Correlation measurements (relating to Figure 6). (A) TSDN spectral tuning raw data (dashed grey), smoothed data (solid black, filter window 191 nm), minima (dashed red), maxima (dashed magenta), and half-max (dashed blue). (B) Spectral tuning first and second derivatives (solid black, dashed grey), and first derivative zero crossings (dashed green). (C) Comparison of raw TSDN data (dashed grey) with smoothened TSDN spectral tuning (solid colors) using different filter window sizes. Filter window of 191 used for all calculations. (D) Cosine similarity between TSDN spectral tuning and wing reflectance (solid line) and daylight reflection (dashed line) for different smoothing window sizes. (E) Percent difference in cosine similarities between TSDN*Reflectance and TSDN*Daylight.

